# Extensive diversity and rapid turnover of phage defense repertoires in cheese-associated bacterial communities

**DOI:** 10.1101/2022.04.13.488139

**Authors:** Vincent Somerville, Thibault Schowing, Hélène Chabas, Remo S. Schmidt, Ueli von Ah, Rémy Bruggmann, Philipp Engel

**Author notes:** Corresponding author, Department of Fundamental Microbiology, Biophore, University of Lausanne, 1015 Lausanne, Switzerland. These authors contributed equally (Vincent Somerville, Thibault Schowing).

## Abstract

Phages are key drivers of genomic diversity in bacterial populations as they impose strong selective pressure on the evolution of bacterial defense mechanisms across closely related strains. The pan-immunity model suggests that such diversity is maintained because the effective immune system of a bacterial species is the one distributed across all strains present in the community. However, only few studies have analyzed the distribution of bacterial defense systems at the community-level, mostly focusing on CRISPR and comparing samples from complex environments. Here, we studied 2’778 bacterial genomes and 158 metagenomes from cheese-associated communities, which are dominated by a few bacterial taxa and occur in relatively stable environments. We find that nearly identical strains of cheese-associated bacteria contain diverse and highly variable arsenals of innate and adaptive (i.e CRISPR-Cas) immunity suggesting rapid turnover of defense mechanisms in these communities. CRISPR spacer abundance correlated with the abundance of matching target sequences across the metagenomes providing evidence that the identified defense repertoires are functional and under selection. While these characteristics align with the pan-immunity model, the detected CRISPR spacers only covered a subset of the phages previously identified in cheese, suggesting that CRISPR does not provide complete immunity against all phages, and that the innate immune mechanisms may have complementary roles. Our findings show that the evolution of bacterial defense mechanisms is a highly dynamic process and highlight that experimentally tractable, low complexity communities such as those found in cheese, can help to understand ecological and molecular processes underlying phage-defense system relationships.

**Importance:** Bacteria are constantly exposed to phage predation and hence harbor highly diverse defense arsenals. According to the pan-immunity hypothesis the effective immune system of a bacterial species is not the one encoded in a single genome but in the entire community. However, few studies have investigated how defense systems are distributed within communities. Here, we carried out (meta)genomic analyses of bacterial communities used in cheesemaking. These are tractable communities of biotechnological interest which house few bacterial species and are exposed to high phage pressure. In line with the pan-immunity hypothesis, we find that nearly identical strains of cheese-associated bacteria contain highly variable arsenals of innate and adaptive immunity. We provide evidence for the functional importance of this diversity, and reveal that CRISPR alone does not provide complete immunity against all phages. Our findings can have implications for the design of robust synthetic communities used in biotechnology and the food industry.

## Introduction

Bacteria have evolved diverse defense systems to cope with the parasitic lifestyle of phages (1, 2). These systems can be divided into the innate and the adaptive “prokaryotic immune system” (3). Classical examples of innate immunity are restriction-modification (4) or abortive infection systems (5). However, many additional innate immune mechanisms have recently been discovered highlighting the strong selective pressure imposed by phages on microbial communities (2, 6). The only ‘adaptive’ immune system known so far is the CRISPR-Cas system (Clustered Regularly Interspaced Short Palindromic Repeat - CRISPR Associated). It is based on the incorporation of short DNA sequences of phages or other genetic elements (so-called spacers) into dedicated CRISPR arrays encoded in the bacterial genome. Upon a phage encounter, the transcribed spacers bind to the phage DNA and target it for degradation via the Cas proteins (7).

Phage defense systems are prevalent across bacteria and most bacteria harbor several systems in their genome (2, 8). However, their distribution varies across bacteria. For example, CRISPR-Cas systems are found in 40% of all known bacterial species (9), while Viperins are restricted to a few taxonomic groups (10). Moreover, phage defense systems are often strain-specific (11), i.e. closely related bacteria can harbor completely different arsenals of defense systems. Various factors have been discussed to influence the distribution of defense systems across bacteria (12). Notably, the presence of CRISPR-Cas has been associated with environmental factors such as temperature, oxygen levels or phage abundance (13–15). Also, genetic incompatibilities of defense systems with other cellular functions, including other defense systems, have been reported (16, 17).

Innate immune systems often cluster in genomic islands and are associated with mobile genetic elements (18–20) suggesting an important role of horizontal gene transfer (HGT) for defense system evolution. The ecological relevance of such genomic plasticity was demonstrated by a recent study which showed that nearly clonal isolates of *Vibrio sp*. are resistant to diverse phages due to the presence of distinctive defense islands acquired via HGT (21).

Based on the observation that phage defense mechanisms show a high extent of genetic turn over, the pan-immunity hypothesis has been proposed, which states that the effective immune system of a bacterial species is not the one encoded in a single genome, but in the pan-genome of the entire population (22). In other words, while a single strain cannot carry all possible defense systems, the presence of nearly clonal strains with different defense systems increases the available arsenal of defensive mechanisms via HGT and thus increases the resistance of the entire population (pan-immunity).

Comparative genomics of closely related isolates combined with shotgun metagenomics provide excellent opportunities to assess the diversity of phage defense systems in natural bacterial communities and can help to advance our understanding of their evolutionary ecology (23). Two recent shotgun metagenomic studies have looked at CRISPR spacer diversity in microbial communities, one focusing on environmental samples from the Earth Microbiome Project (15) and another one focusing on diverse samples from the human microbiome (24). Meaden et al. identified a positive association between CRISPR spacer and the abundance of the corresponding phage target sequences (protospacer), suggesting that there is a direct link between phage pressure and the maintenance of corresponding spacer sequences. Münch et al. identified differences in the prevalence of CRISPR spacers between different human body sites suggesting the existence of niche-specific phage populations. However, none of the two studies looked at the diversity of innate immune systems. Moreover, in both studies highly diverse communities from heterogeneous environments were compared providing limited insights into the intraspecific variation of phage defense systems and their evolutionary turnover in bacterial populations.

Here, we focused on cheese-associated bacterial communities. These communities harbor only a few bacterial species, have been propagated in relatively stable environments (i.e. cheese or milk) over generations, and are known to be exposed to diverse phages (25–28). This makes them tractable systems to study the evolution of phage defense systems in microbial communities at the strain-level (29, 30). Indeed, our previous study of a single Swiss cheese starter culture has shown that extensive intraspecific CRISPR spacer diversity exists in these otherwise nearly clonal populations of bacteria (31).

Here, we expanded this analysis to all publicly available genomic datasets from cheese-associated communities (excluding cheese rind) comprising 26 bacterial species, 2’778 genomes, and 158 metagenomes. We determined the distribution of both innate and adaptive immune systems across these datasets and quantified the diversity, abundance, turnover rate, and viral targets of all CRISPR spacers. We find that (i) cheese-associated bacteria contain an unprecedented high degree of diversity in phage resistance mechanisms across nearly identical strains, (ii) there is a strong correlation between CRISPR spacer and phage abundance, and (iii) CRISPR spacers only provide immunity to a subset of the phages identified in cheese. These results indicate highly dynamic bacteria-phage interactions driving genomic plasticity in cheese-associated environments and are compatible with the pan-immunity model of the evolutionary ecology of defense systems in microbial communities.

## Results and Discussion

### Strain-specific immune gene arsenals across cheese-associated bacteria

In order to obtain an overview of the diversity of phage defense systems in cheese-associated bacteria, we first determined which taxonomic groups are prevalent across cheese-associated communities by profiling 480 community samples from 18 different studies (Suppl. table 1). This included 322 16S rRNA gene amplicon sequencing and 158 shotgun metagenomics datasets from both mesophilic (cooked at lower temperatures) and thermophilic (cooked at higher temperatures) cheese starter cultures and ripened cheese (Fig. 1A/B). We excluded cheese rind samples from this analysis, because they consist of highly variable microbial communities with a more complex ecology (32). A total of 196 species were identified of which 26 were present in >2% of all samples with a median abundance of >0.1% (Fig. 1B and Suppl. Fig. 1). The large majority of these species were from the order *Lactobacillales* with the two major species of mesophilic and thermophilic cheese samples, *Lactococcus lactis* and *Streptococcus thermophilus*, dominating the communities (Suppl. Fig. 2).

**Figure 1.**
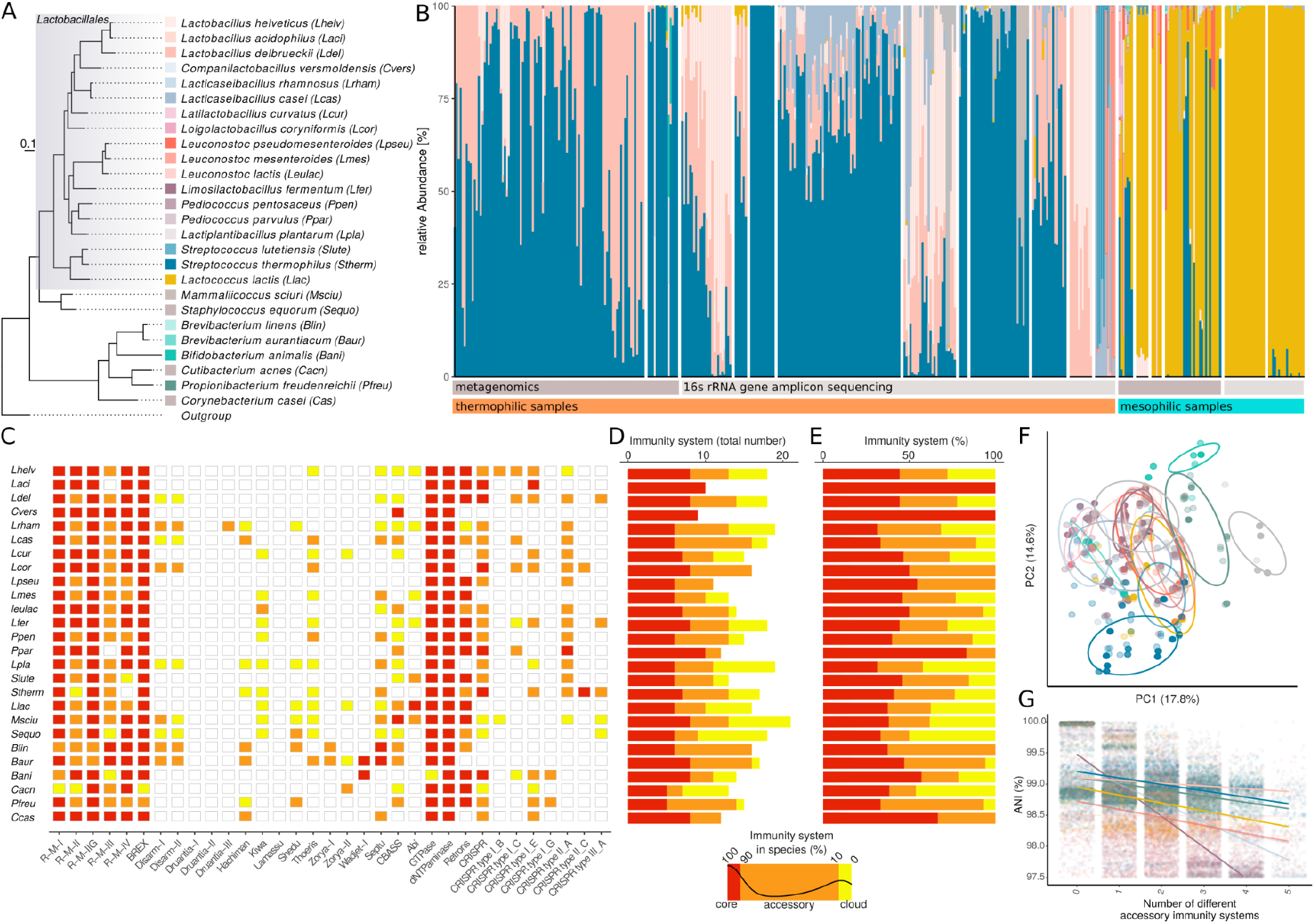
Diversity of phage defense systems in the genomes of cheese-associated bacterial species. A) Core genome phylogeny of the 26 predominant species found in the cheese-associated communities and their corresponding color key used in Figure 1A. B) Species-level composition of cheese-associated communities (starter and non-starter) grouped by studies. Sample type and community profiling method (16S rRNA gene amplicon or shotgun metagenomics sequencing) is indicated. C) Heatmap illustrating the fraction of genomes per species containing different innate immunity mechanisms and CRISPR and CRISPR subtypes; The color scheme is indicated below D) and E). D) The absolute and E) relative count of core (>90% of strains), accessory (90%< of strains >10%) and cloud (<10% of strains) defense systems. F) Principal component analysis of all strains based on the abundance/presence of different defense systems (colored according to legend in A). G) The number of different defense systems vs. average nucleotide identity between two genomes of the same species. Including only the most dominant species comparisons. The statistics of the regression lines are illustrated in Suppl. Fig. 4 and the colors corresponds to the legend in A)

We next retrieved 2’778 genomes of the 26 predominant species from NCBI and the *in house* genomic database of Agroscope (Suppl. Fig. 1). The genomes were screened for the presence of homologs of 25 different phage defense systems using a hmm-search approach. In total, 17’565 innate immune systems and 1’972 CRISPR-Cas systems were identified. Restriction/Modification (RM) systems (Fig. 1C-E), GTPases, deaminases, and retrons were common innate immune systems across almost all species. On the contrary, only a few species harbored homologs of e.g. Abi systems (Suppl. Fig. 3). All species contained CRISPR-Cas systems with the exception of *Brevibacterium aurantiacum, Brevibacterium linens, Companilactobacillus versmoldensis, Lactococcus lactis* and *Leuconostoc mesenteroides* (Fig. 1C). None of the defense systems were found to be specific to a given species. Moreover, species did not cluster by defense systems composition (Fig. 1F) suggesting overlapping defense strategies across species. On average we found 7.5 (sd=1.4, Fig. 1D) defense systems per genome with all species harboring more defense systems than previously described for bacteria of other environments (11). Considering that the number of defense systems reflects the extent of phage pressure in a given environment, this supports the idea that phages are prevalent in cheese-associated communities (11).

Notably, only 49% (sd=19%) of the defense systems detected within a species were shared among all strains of that species (core), 35% (sd=14%) were shared between 10-90% of the strains (accessory), and 25% (sd=13%) were present in less than 10% of the strains (cloud) (Fig. 1D/E). Similarly, while CRISPR-Cas systems were detected in almost all species, only 49% (sd= 40%) of the strains of these species encoded CRISPR-Cas systems in their genome (Fig. 1E). The number of shared innate immune systems decreased with increasing genomic divergence as measured by pairwise Average Nucleotide Identity (ANI) (Fig. 1G, Suppl. Fig. 4).

No correlation was found between the presence of different innate immune systems across the analyzed genomes (Suppl. Fig. 5). However, we did find that species without CRISPR-Cas systems harbored significantly more innate defense mechanisms than species with CRISPR-Cas (unpaired Wilcoxon test, p-value<0.001, Suppl. Fig. 6). Interestingly, this pattern was reversed when comparing strains of the same species (Wilcoxon test, p-value<0.001, Suppl. Fig. 6); i.e. strains without CRISPR tended to have fewer innate defense mechanisms than strains with CRISPR. As phage defense systems are costly to maintain, the loss of such genes could be the result of extensive passaging of certain strains in phage-deprived environments, especially as many of the sequenced genomes come from laboratory strains.

### Rapid turnover of CRISPR spacers in nearly identical strains of cheese-associated bacteria

To assess if the high diversity of the innate immune system is paralleled by a similarly high extent of diversity in adaptive immunity, we identified all CRISPR spacers across all 2’778 genomes of the 26 predominant cheese-associated bacterial species. We detected a total of 1’972 CRISPR arrays containing 16’506 unique spacers (Suppl. Fig. 7). The number of spacers per array (median = 25.83, sd = 19.29) varied within and across species (Suppl. Fig. 7) and also between the different CRISPR-Cas subtypes detected in the analyzed datasets (Suppl. Fig. 8).

As expected, no spacers were shared between genomes belonging to different species or between arrays from different CRISPR-Cas subtypes. However, also within species (ANI>95%), only 41% of all genome pairs shared spacers. Moreover, the fraction of shared spacers between two genomes was relatively small (median = 9.2%, sd = 33.8%). Even nearly identical strains with an ANI >99.5% shared on median as little as 55% of their spacers (sd=27%). The proportion of shared spacers decreased with genomic distance (as measured by decreasing pairwise ANI) (slope=41.3,R^2^=0.48, Fig. 2A) in all species (Suppl. Fig. 9). This is in line with the observed decrease in shared innate immune systems with increasing genetic distance between strains. Although there seems to be a signature of vertical evolution over very short evolutionary timescales, the results overall suggest that most spacers are not maintained for very long but are continuously gained and lost. The only exception concerns a subset of divergent *L. casei* genomes (ANI∼98%, Fig. 2A), which contained plasmids carrying a CRISPR array sharing >25% of the spacers.

**Figure 2.**
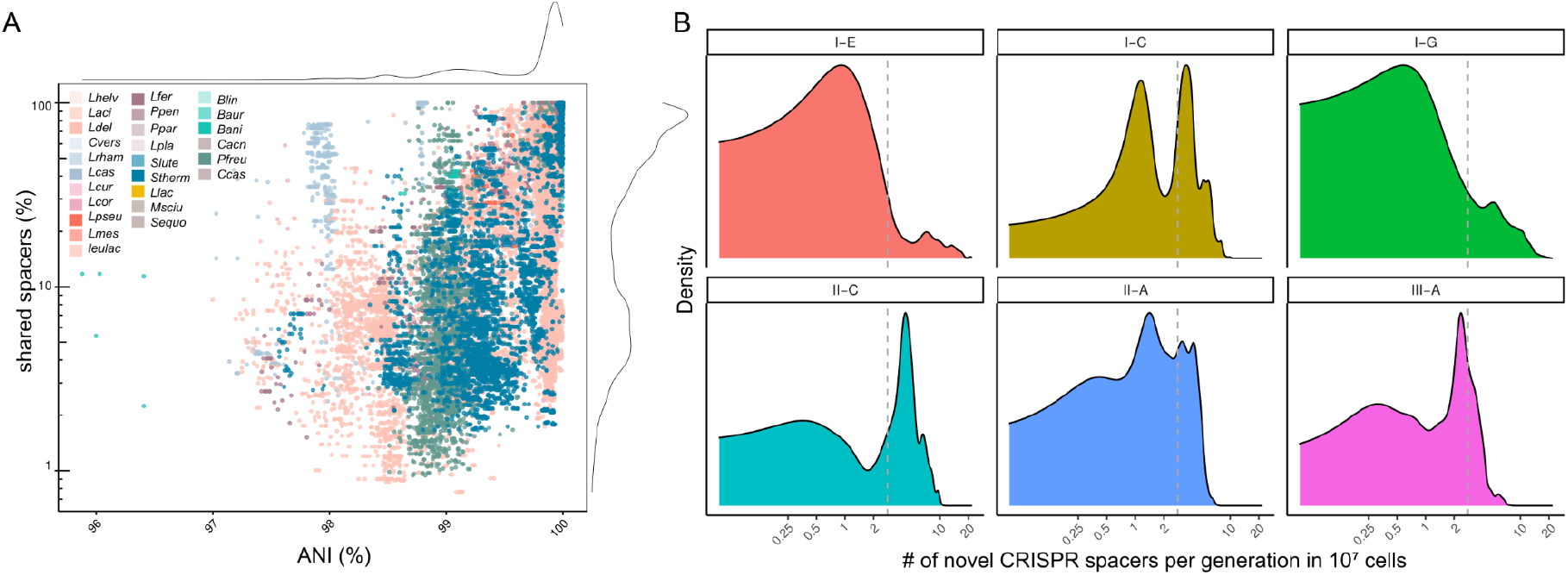
High turnover of CRISPR spacers in cheese-associated bacterial genomes. A) Pairwise comparison of the average nucleotide identity (ANI) and the fraction of shared spacers between genomes of the same species (n=160’556 comparisons). B) Density plots of the number of novel CRISPR spacers acquired per generation in a microbial community of 10^7^ cells subdivided into the six different CRISPR-cas subtypes. The dashed line indicates the median spacer turn-over rate.

To obtain an estimate of the CRISPR spacer turnover rate, we calculated how many novel CRISPR spacers would be acquired in each new generation in a community of defined size. We divided the number of unique spacers per genome by the number of nucleotide differences for each genome pair with an ANI >99%. We found that one unique CRISPR spacer corresponds on average to 1’355 core genome single nucleotide variants (SNVs) (sd=2492). When accounting for a core genome mutation rate of 8.9 × 10^−11^ mutation per base-pair per generation (33) and a cell density of 10^7^ cells per community, we calculated that the CRISPR turnover rate corresponds to a median of 2.8 CRISPR spacers per generation (Fig. 2B). This suggests that the acquisition of novel CRISPR spacers is extremely rapid and that at every bacterial generation several novel spacers can be incorporated. However, as we only considered fixed mutations, we may underestimate the time of divergence between these genomes which would result in a lower turnover rate. Indeed, previous estimates of CRISPR acquisition rates based on experimental data were ∼1 magnitude lower (<0.1 spacer/generation in (31) or ∼0.5 spacer/generation in (34)).

Interestingly, we observed marked differences in spacer turnover rates between different CRISPR-Cas subtypes but not between different species. More specifically, the spacer turnover rates of arrays belonging to CRISPR-Cas subtypes I-E, I-G and III-A were generally lower than the median turnover rate (2.8 spacers per generation). On the contrary, spacers turnover rate of the CRISPR-Cas subtype II-C was generally higher than the median turnover rate. Finally, the turnover rates of arrays belonging to CRISPR-Cas subtypes I-C and II-A showed a bimodal distribution with some arrays having high and others low rates of spacer turnover (Fig. 2B). Variation in spacer turnover rate has been previously observed and was suggested to reflect differences in phage pressure acting on the different strains (35). Our data suggest that it also depends, at least partially, on intrinsic properties of the CRISPR-Cas subtype within the species (Suppl. Fig. 10).

### Extensive CRISPR spacer diversity in metagenomic datasets of cheese-associated communities

The high turnover rates of CRISPR spacers estimated from the isolate genomes suggests the presence of high levels of CRISPR spacer diversity within and across cheese-associated communities. Based on the identification of flanking CRISPR repeats we extracted 8’226 non-redundant full-length spacer sequences from the Illumina reads of the 158 shotgun metagenomic samples presented in Figure 1B. On average 5.24 (sd=6.23) spacers per million reads were identified per sample (Fig. 3A). There was no difference in CRISPR spacer diversity between mesophilic (cheese that is made at ∼30 °C) and thermophilic (cheese that is made at >45 °C) communities. This was surprising, as mesophilic cheese communities are dominated by the non-CRISPR containing species *L. lactis* and *Leuc. mesenteroides*, and suggests that subdominant community members harbor a high number of CRISPR spacers.

**Figure 3.**
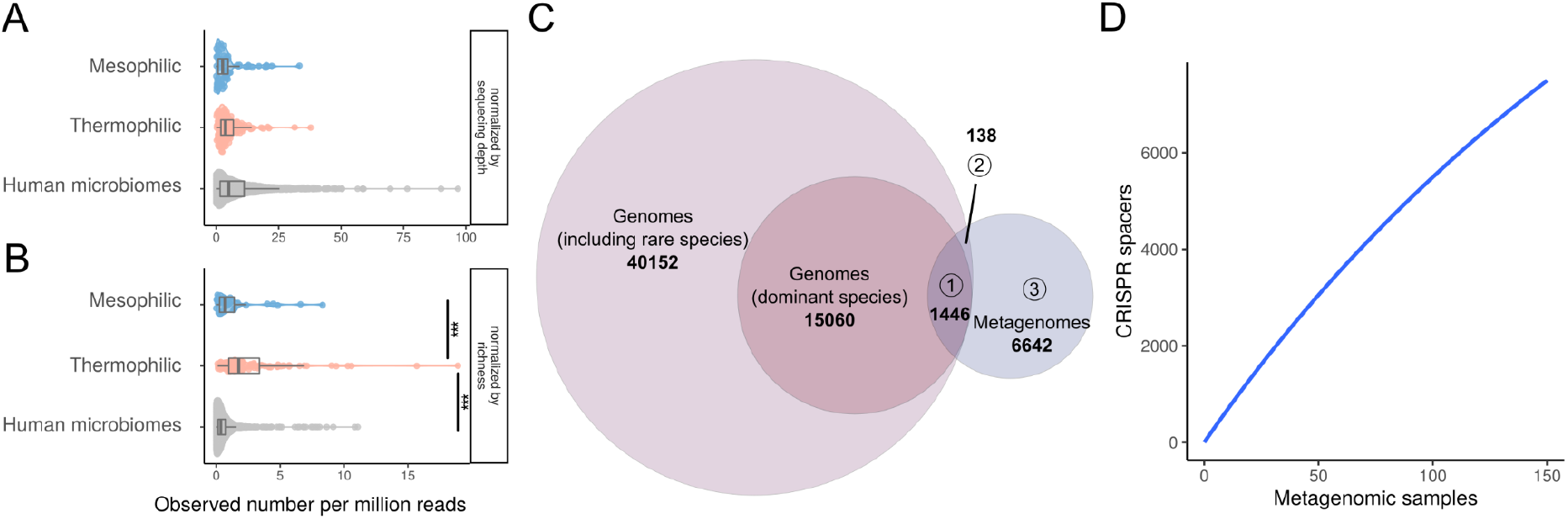
Metagenomic CRISPR diversity. A-B) Number of CRISPR spacers present in the different metagenomic samples normalized by A) the sequencing depth and B) the sequencing depth and the species richness. (*** illustrates Wilcoxon p-values < 0.001). C) The number of spacers detected in the isolated genomes of predominant and subdominant cheese community species and in the shotgun metagenomic samples. Intersection of circles shows the number of shared CRISPR spacers (intersection(1)=metagenomic and dominant species, intersection(2)=metagenomic and subdominant species, intersection(13=only metagenomic). D) The cumulative plot (rarefaction curve) of the CRISPR spacers detected in the metagenomic samples.

We compared our dataset to a previously published analysis of CRISPR spacer diversity in human microbiomes and found that the diversity of CRISPR spacers in cheese-associated communities and the human microbiomes is not significantly different from each other (Fig. 3A). However, when accounting for the higher species diversity in the human microbiome (i.e. by normalizing to the richness of each community), we found that thermophilic communities harbor more CRISPR spacers per species than human microbiomes (Fig. 3B). This is in line with previous studies, which had shown that high CRISPR-Cas diversity is associated with anaerobic growth, high temperatures and non-host environments (13–15), all of which are characteristics of cheese environments.

Surprisingly, only a small fraction of the spacers identified across the metagenomic datasets (17.6%) matched to spacers detected in the 2’778 isolate genomes of the predominant cheese-associated species (Fig. 3C). When also considering genomes of subdominant species, this number increased only little to 1’584 (19.3%) matching CRISPR spacers. As no other species were detected in the analyzed metagenomes (Fig. 1A/B), we conclude that the large majority (80.7%) of the metagenomic CRISPR spacers corresponds to within-species diversity not covered by the currently available isolate genomes. Further, we found little overlap in CRISPR spacer diversity between metagenomes. Only very few spacers (8%) were shared amongst more than two metagenomes (Suppl. Fig. 11). Moreover, a rarefaction curve analysis showed that with the addition of each metagenomic sample new spacers are being discovered (Fig. 3D). Together, this indicates that the CRISPR spacer diversity is extensive in cheese-associated communities and that we have only detected a fraction of this diversity in our study.

### CRISPR spacer abundances correlates with target abundance

If the spacers have any ecological relevance (36, 37), one would expect to find a positive correlation between the abundances of phages and their matching spacer sequences. To quantify both spacer and phage (i.e. target) abundance directly from the metagenomic samples we mapped the metagenomic reads of the 158 datasets to all dereplicated repeat-spacer-repeat sequences identified in the 2’778 isolate genomes. As spacers are usually shorter than Illumina reads, reads containing spacer and repeat sequence were considered to come from a CRISPR array (hereafter referred to as spacer reads). In contrast, reads mapping to only spacer sequences were considered to come from a target (e.g. a phage, hereafter referred to as protospacer reads) (Suppl. Fig. 12).

In each metagenome, we identified between 41 and 1’961 repeat-spacer-repeat sequences, which recruited at least one spacer or protospacer read. In many cases only protospacers (41.3%) or spacer (33.6%) reads were identified. For the remaining 25.1% of cases, we found a positive correlation between spacer and matching protospacer abundance (slope=0.52, R^2^=0.37, Fig. 4A), independent of the CRISPR-Cas subtype or the metagenomic sample (Suppl. Fig. 12/13). This is in line with previous results obtained for the Earth Microbiome Project (15) and supports the idea that spacers targeting highly abundant phages are under positive selection and thus dominant in the community (38). Notably, in our previous study focusing on a single cheese starter culture we had found the opposite pattern. This may be explained by the fact that a single phage dominated this community, causing chronic infections and thereby overcoming CRISPR-based immunity (31). Interestingly, with increasing spacer abundance the ratio of protospacer to spacer abundance decreased (slope=-0.48, R^2^=0.32, Fig. 4B), suggesting that highly abundant spacers are effective in decreasing protospacer abundance. A similar correlation has previously been described for the viral fraction outside of the cells measured by virus-microbe-ratios (VMR) (39, 40).

**Figure 4.**
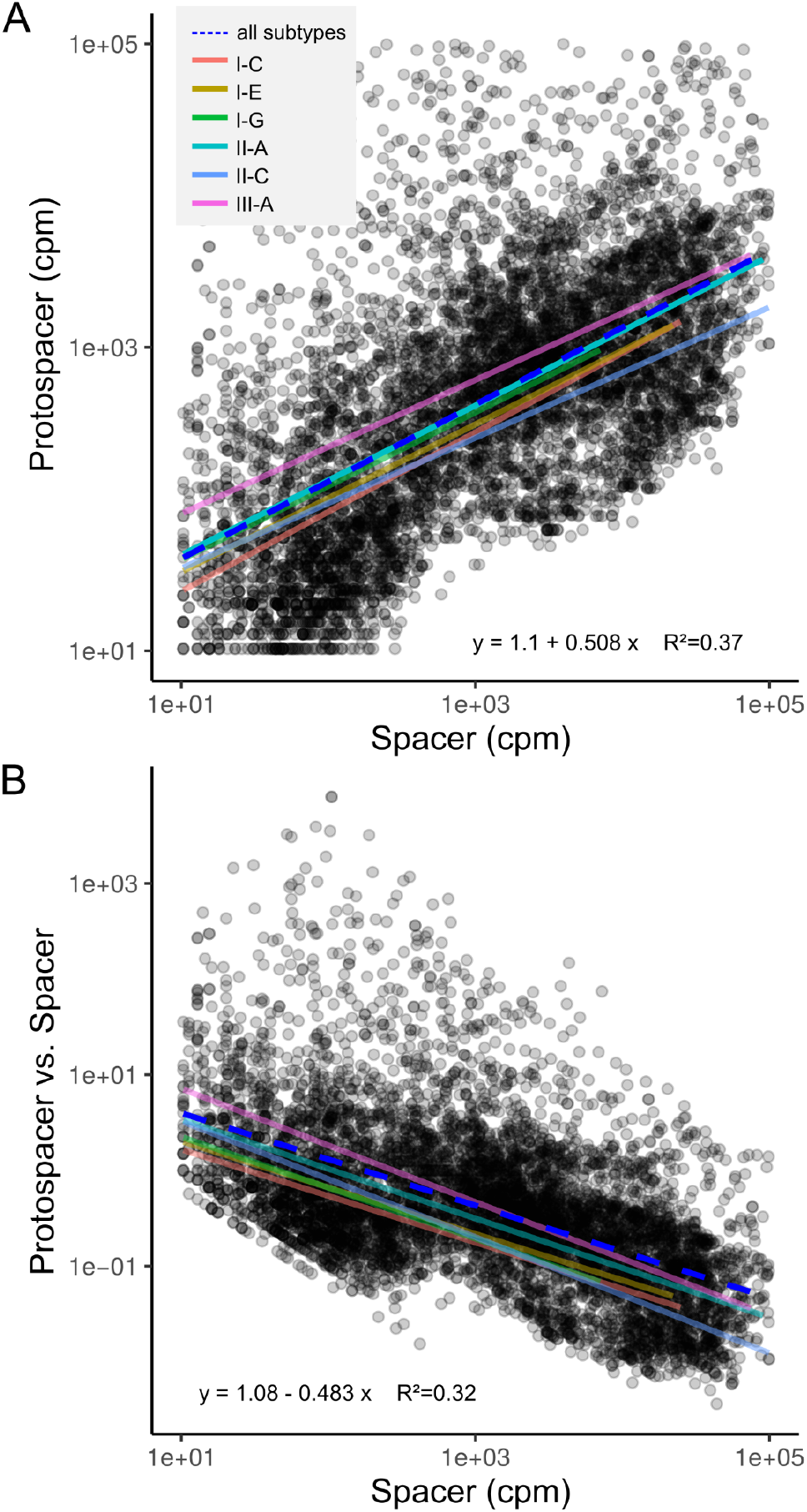
Metagenomic CRISPR spacer and protospacer abundance. A) The protospacer and spacer abundance of all metagenomic samples indicated in counts per million (cpm). The dashed blue and colored lines indicate the linear regression across all and specific CRISPR-Cas subtypes, respectively. The correlation values relate to all subtypes. B) The spacer abundance in relation to the ratio of protospacer versus spacer abundance.

Further, we wanted to see if the CRISPR spacers are genetically linked and verify if the entire strain or the individual spacers are the level of selection. Therefore we looked at the spacer abundances within one metagenomic sample. The spacer abundance within a single metagenome clearly has a non-normal distribution, with few spacers being abundant, while the majority are of low frequency (Suppl. Fig. 14). This indicates that individual spacers can sweep through the population rather than the entire strain being selected for.

### No complete pan-immunity by CRISPR defense

For a large number of the spacers identified in the isolate genomes, we did not find a corresponding protospacer sequence in the metagenomes. However, in 41.3% of the cases (see above) we identified a protospacer, suggesting that not for all targets present in a given sample, CRISPR spacers have been acquired at a detectable level. To determine the identity of sequences targeted by the CRISPR-Cas system, we searched the 63’438 CRISPR spacers identified in the 2’778 genomes and 158 metagenomes against the NCBI nucleotide database (i.e. bacteria), the IMG viral database (i.e. phages), and the plasmid PLSDB database (41). In total 49% of the CRISPR spacers had a hit (maximal 2 mismatches) in at least one of these databases. The majority of spacers matched to phages (36%) followed by bacterial chromosomes (9%) and plasmids (3%) (Fig. 5A). These proportions are in a similar range to what has previously been reported for other bacteria (42). Most of the sequences matching bacterial chromosomes and plasmid were found in genes belonging to the COG (Clusters of Orthologous Groups) category “Replication and repair”, which includes selfish elements such as transposons (Suppl. Fig. 15). The fraction of spacers targeting phages varied across species. In case of *S. thermophilus*, 61% of the spacers mapped to known phages, while for other species, e.g. *L. delbrueckii* or *L. helveticus*, only 30% and 36%, respectively, mapped to known phages, probably reflecting the extent of research conducted on the phage diversity of different bacterial species.

**Figure 5.**
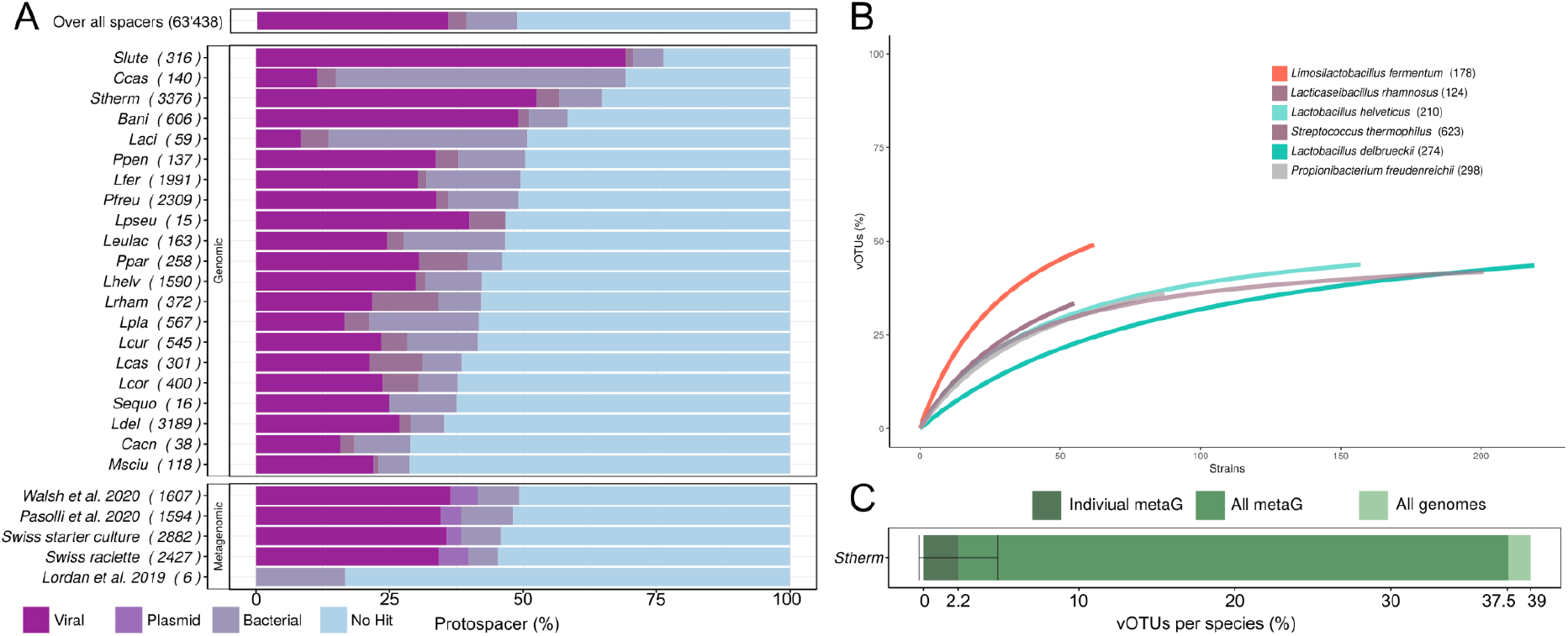
Protospacer diversity. A) The fraction of CRISPR spacers mapping to the Viral IMG db, the bacterial NCBI database or having no map. Each species (top) and metagenomic project (bottom) are subdivided and the number of spacers therein are indicated in the brackets. B) The rarefaction curves of vOTU for all species with more than 50 genomes and more than 85 described vOTUs. C) The fraction of IMG vOTUs targeted by all metagenomic samples (green bar), one metagenomic sample (dark green bar) or all metagenomic samples in that project (light green bar) of *S. thermophilus* (total 623 vOTUs). The bars indicate the standard error.

**Figure 6.**
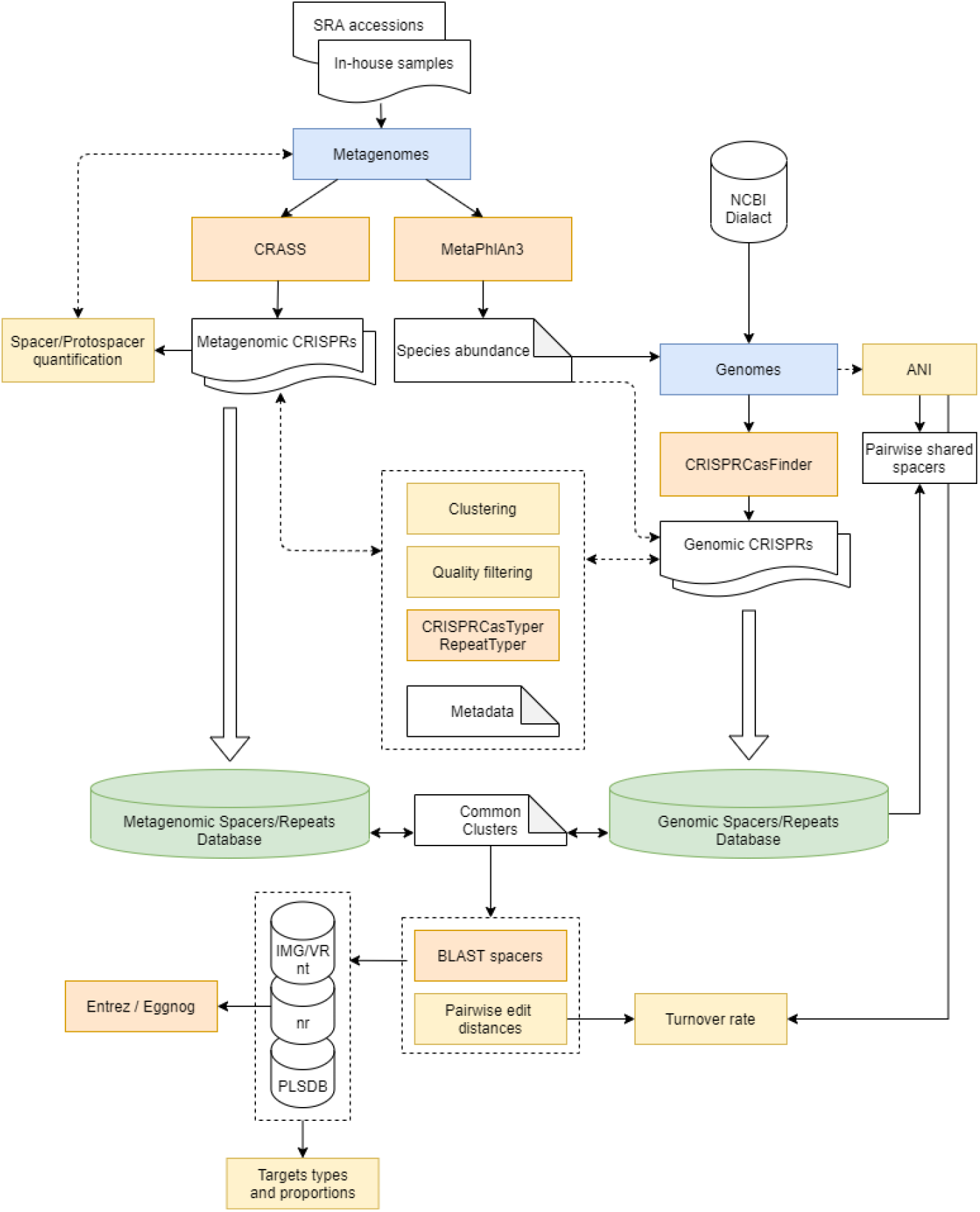
Overview of the computational workflow. Main tools are represented in orange, main operations in yellow and the two main datasets in green.

To assess the range of phages targeted by a given isolate, we categorize the phages into discrete viral Operational Taxonomic Units (vOTU) inferred by the viral IMG database (43). Further, we limited this analysis to the genomes of the six bacterial species with the largest diversity of phages represented in the database (i.e. > 85 described vOTU), namely phages of *S. thermophilus, L. delbrueckii, L. helveticus, L. fermentum, L. rhamnosus* and *P. freudenreichii*. Within these species, we observed that each strain targets on average 9 vOTUs (sd=8) (Suppl. Fig. 16), which corresponds to 2.5% (sd=0.6%) of the known phage diversity of these species. This is in line with a recent study where they modeled that 1-10% of the total phage diversity is covered by the CRISPR-Cas system of a single strain (44). Notably, most isolate genomes (83%) harbored only a single spacer against a given vOTU (Suppl. Fig. 17) with no major differences between species (Suppl. Fig. 18). This observation is in contrast to what has been observed in laboratory studies of phage-bacteria coevolution (36, 45) or in cases of chronic phage infections in *S. thermophilus* (31), where multiple spacers targeting the same vOTU were often found to be integrated into the CRISPR array.

Our results seem to suggest that the spacer repertoire of a given strain is aimed at targeting a broad range of different phages rather than being specialized towards a single vOTU. To assess whether the presence of several strains with diverse CRISPR spacers provides pan-immunity against a broad range of phages, we conducted a rarefaction analysis of the CRISPR-vOTU matches identified in the isolate genomes. For all six analyzed species, the curve rapidly flattened with no more than 50% of the known vOTU having matching CRISPR spacers in the analyzed isolate genomes (Fig. 5B). For example for *S. thermophilus* only 39% of all phage vOTUs had matching spacers across the isolate genomes. This indicates that no combination of isolates results in complete CRISPR-based immunity against all known phages of a given species. Analysis of the metagenomic CRISPR spacers mapping to *S. thermophilus* phages confirmed these results: only 37.6% of the known *S. thermophilus* phages were targeted by CRISPR spacers identified across the 158 metagenomic datasets (Fig. 5C). Even fewer phages were targeted by spacers identified in individual metagenomic datasets (mean=2.5%, sd=2.2, Fig. 5C). Phages not targeted by any spacer did not seem to be rare as the vOTU clusters were not necessarily smaller than the clusters of targeted vOTU (Suppl. Fig. 19). Moreover they did not contain more anti-CRISPR genes than phages that had matching CRISPR spacers in the communities (Suppl. Fig. 20). It is possible that these phages are integrated as prophages in the bacterial genomes and thereby avoid CRISPR-based immunity or that the bacteria and phages have not encountered each other due to spatial population structure or segregation into different communities that have not yet been sampled (46).

## Conclusion

Previous studies have used shotgun metagenomics to characterize CRISPR diversity in bacteria found on the human body, in the ocean or the soil (15, 24). Cheese-associated communities are much simpler than these previously analyzed communities (47, 48). They contain much fewer species and are propagated in relatively stable environments (49, 50). A large amount of genomic data is available for cheese-associated communities as they are established experimental model systems to study bacteria-phage interactions. This allowed us to assess the intraspecific diversity and evolutionary dynamics of phage defense systems across a wide range of bacterial species and communities by analyzing publicly available genomes and metagenomic datasets. We found extensive diversity in innate and adaptive immune defense mechanisms across cheese-associated bacteria, despite the overall little genomic diversity present in these communities. Phages are known to be common in these environments and pose a risk for the cheese making process (28). However, the extent of defense systems we have found seems to exceed the diversity present in previously analyzed ecosystems, which is surprising given that cheese communities are closed systems with little opportunities for migration/invasion.

Our analysis revealed that innate immune systems were distributed in a strain-specific manner across the analyzed genomes of cheese-associated communities. Likewise, CRISPR spacer repertoires varied substantially across nearly clonal isolates and the amount of CRISPR spacers present in the metagenomic datasets seemed infinite. Accordingly, our estimation of the CRISPR spacer turnover rates suggested rapid gain and loss of CRISPR spacers with important differences found between different CRISPR-Cas subtypes but not necessarily species. The ecological relevance of the identified diversity is highlighted by the finding that metagenomic CRISPR spacers matching abundant target sequences (e.g. phages) were also abundant in the corresponding metagenomic sample which indicates specific bacteria-phage responses in terms of their ecological dynamics.

Our observations align with the pan-immunity model proposed for understanding the evolutionary ecology of innate immune systems (22) and suggest that this model can be extended to CRISPR-based adaptive immunity, as the three key points of this model seem to be fulfilled. First, we show that there is standing genetic diversity in CRISPR spacer diversity in cheese-associated bacteria and communities. Second, similar as for the horizontal mode of transfer proposed for the innate immune systems (21), the high turnover of spacers detected in our study reflects that novel defensive repertoires can be rapidly acquired (and lost) in the population. Third, CRISPR spacers seem to be selected for and hence functional as their abundance correlates with phage abundance.

On the contrary, for about 50% of the phages that were isolated from cheese-associated communities, we could not identify any matching CRISPR spacers across the genomes/metagenomic datasets. This may hint at the combined importance of both innate and adaptive immune systems and could suggest that CRISPR is not effective against all phages. It is possible that these phages avoid CRISPR targeting, making them interesting candidates to look for novel anti-CRISPRs mechanisms. Overall, these discoveries allow us to not only to understand the evolutionary ecology of phage-bacteria interactions but could also be instrumental in improving the protection of cheese starter culture by developing phage-based therapy (51) as a protection against pathogens (52) or invasive strains (53).

## Materials and methods

### Metagenomes

In order to analyze species and CRISPR diversity in cheese related samples we gathered 158 shotgun metagenomic samples and 322 16S metagenomic samples from overall 18 studies from NCBI (see Suppl. table 1) (31, 54–68).

### Metagenomic species profiling

To determine the species composition of the metagenomic samples, we used MetaPhlAn (v3.0.7-1) (69). For the 16S rRNA gene amplicon sequencing datasets we used the extensive FoodMicrobionet (8.2020) collection and analysis pipeline (see script section) (70, 71). Overall, 185 species were identified. We selected the dominant species which were present in >2% of samples and had a median overall abundance >0.1%. From those species we randomly selected one genome and created a species tree with BCGtree (v1.1.0) (72).

### Genomes analysis

All genomes (or max. 500 newest genomes if more were available) for the 185 identified species were downloaded from NCBI RefSeq (17.05.2021). For the 26 dominant species, we additionally integrated all available genomes from the *in-house* cheese database of Agroscope called “Dialact”.

### Detection of CRISPR and defense mechanisms

The genome assemblies were annotated with CRISPRCasFinder (v4.2.20-1) (9). The raw JSON outputs were parsed and quality filtered with custom Python and R script (see script section). Only spacers with a high evidence level (>4) and shorter than 50bp were retained. CRISPR-Cas subtypes were assigned with CRISPRCasTyper (v1.4.4) (73). Further the remaining defense mechanisms were annotated with defense mechanisms specific HMM files. For the R-M we used previously described HMMs (74). The search was done with hmm-search (v3.1b2) (75).

### Pairwise strains comparison

Average Nucleotide Identity (ANI) was calculated with fastANI (v1.3) (76) and percentage of shared spacers were computed for each pair of strains within an organism. The proportion of shared spacers was calculated as being the proportion of identical spacer clusters between two strains, divided by all spacers.

Additionally, the number of common spacers, the nucleotide diversity and the turnover rate were computed for all strain pairs as well as for each array. The nucleotide diversity was computed as *bidirectional fragment mapping * fragment length (1kb) / ANI* and the turnover rate as *nucleotide diversity / unique spacers*, unique spacers meaning the number of spacers not shared between the two strains. Further, the CRISPR acquisition rate in microbial communities per generation (i.e. CRISPR turnover rate) was calculated as following (turnover rate* cell density of 10^7^ cells)/ mutation rate. When accounting for a mutation rate of 8.9 × 10−11 per bp per generation (33) and a cell density of 10^7^ cells within a community, we were able to calculate a CRISPR acquisition rate in microbial communities per generation (i.e. CRISPR turnover rate, Fig. 2B).

### Detection of CRISPR arrays in metagenomes

In order to identify CRISPR in the metagenomic samples, the raw metagenomic reads were processed using CRASS (v0.3.12) (77) with default parameters. CRISPR spacers, repeats and flanking sequences were then extracted. Each spacer was automatically annotated with coverage information as well as the spacer count per million reads for the whole sample. Spacers with a coverage of 1 were removed as well as spurious spacers of length smaller than 15bp.

### Sequences clustering

Repeats, often referred to as consensus repeats or direct repeats, are not necessarily identical within an array (73) as well as spacers, which do not need a perfect match with their target sequence to be cleaved (78). Thus, repeat and spacer sequences were clustered using CD-HIT-EST (v4.8.1) (79) using a 100% and an 80% identity threshold (24). 80% identity clusters were added in the database separately for genomic repeats, genomic spacers, metagenomic repeats and metagenomic spacers. Venn diagram representations (Fig. 3) have been done with BioVenn (80) and adapted within inkscape, while strictly keeping the calculated surfaces.

### Quantification of spacers and repeats in metagenome

The observed number of spacers and repeats was calculated by dividing the absolute number of spacers and repeats by the total number of reads and later by the total number of bacterial species (richness). Here we also included the data previously described for the human microbiome (24).

### Mapping spacer sequences to reads

In order to quantify the proportion of spacers and protospacers in the metagenomes, we created for every spacer a *repeat-spacer-repeat* sequence and mapped all metagenomic samples individually against this reference. Reads mapping only to the ∼40bp spacer area were assigned to protospacers, whereas reads mapping to the repeat and spacer area were assigned to the CRISPR array. The sum of these counts were normalized by the reads count for each sample.

### Spacers target mapping in viral and bacterial databases

BLASTn (megablast) was performed using an e-value cutoff of 0.01, word-size of 4, keeping only 1 query-subject alignment per pair (max_hsps = 1) and 1 aligned sequence (max_target_seqs = 1) (v2.5.0+). Three databases were used, IMG/VR (v.3.0) (81), nt (non-redundant nucleotide) and the PLSDB plasmid database (v2020_11_19) (41). In order to rule out putative prophage and annotate gene targets, we further blasted to the nr (non-redundant protein) and screened for phage genes and annotated with eggnog (82). Further CRISPR mappings were ruled out by scanning the 2kb up and downstream of the target site for CRISPR repeat annotation in the corresponding genbank files. The assignment to vOTUs was made on the basis of the IMG/VR (v.3.0) (81) mapping.

### Statistics, scripts and data

All statistics were done within R (R Core Team, 2020) and ggplot2 (83). All code for the bioinformatics is available here (https://github.com/ThibaultSchowing/CRISPRscope) and the figures here (https://github.com/Freevini/CRISPRscope). The data is deposited on zenodo (10.5281/zenodo.6444686).

## Acknowledgments

We thank Aline Cuénod, Germán Bonilla-Rosso and Malick Ndiaye for the feedback and discussions of the manuscript. Further we also thank Noam Shani, Petra Lüdin, Verena Schünemann, Abigail Bouwman, Meral Turgay, Marco Meola, Johann Bengtsson-Palme, and the Functional Genomics Center Zurich for the cheese samples sequencing and making their datasets available for this study. The project was funded by Agroscope Switzerland, the University of Lausanne and supported by the Master’s program of the Universities of Fribourg and Bern.

## Author contribution

VS designed and executed the project and wrote the manuscript. TS executed the project and wrote the manuscript. HC, UvA RS and RB advised the project. PE designed the project and wrote the manuscript. All authors gave feedback on the manuscript.

