## Supplemental Information for "Extensive diversity and rapid turnover of phage defense repertoires in cheese-associated bacterial communities"

Vincent Somerville1,2,*,+, Thibault Schowing2,3,+, Hélène Chabas4, Remo S. Schmidt2, Ueli von Ah2, Rémy Bruggmann3, Philipp Engel1
1 University of Lausanne, Switzerland
2 Agroscope, Liebefeld, Switzerland
3 Interfaculty Bioinformatics Unit and Swiss Institute of Bioinformatics, University of Bern, Bern, Switzerland
4 Institute for Integrative Biology, ETH Zürich, Switzerland
+ These authors contributed equally.

### Supplemental Figures:

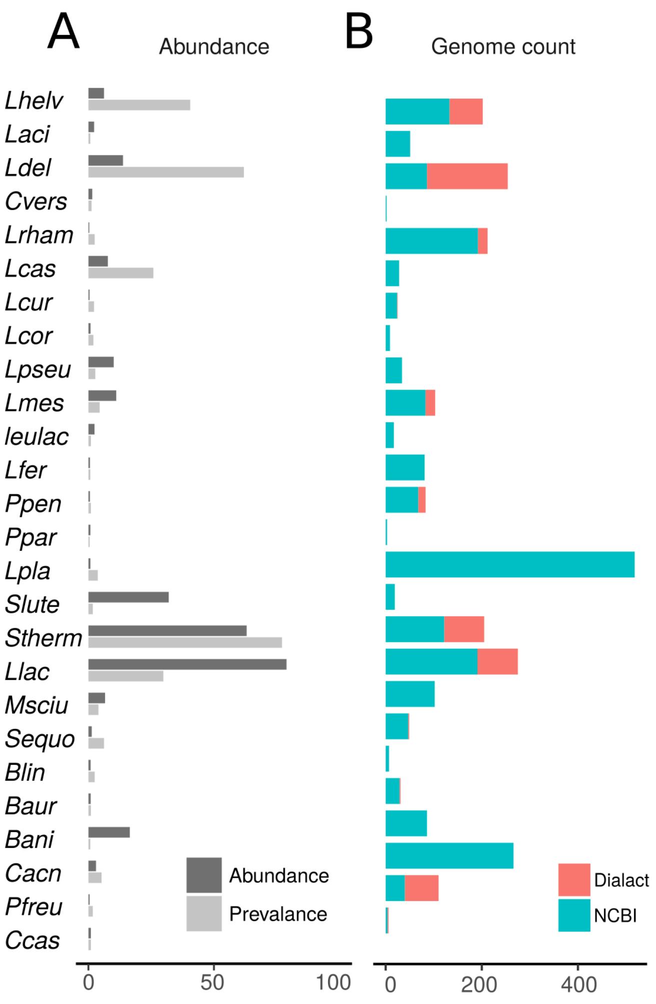


Supplementary Figure 1. A) Mean abundance and prevalence of the different species in the metagenomes illustrated in main Fig. 1A. B). The number of genomes for the different species downloaded from NCBI and in house cheese database.

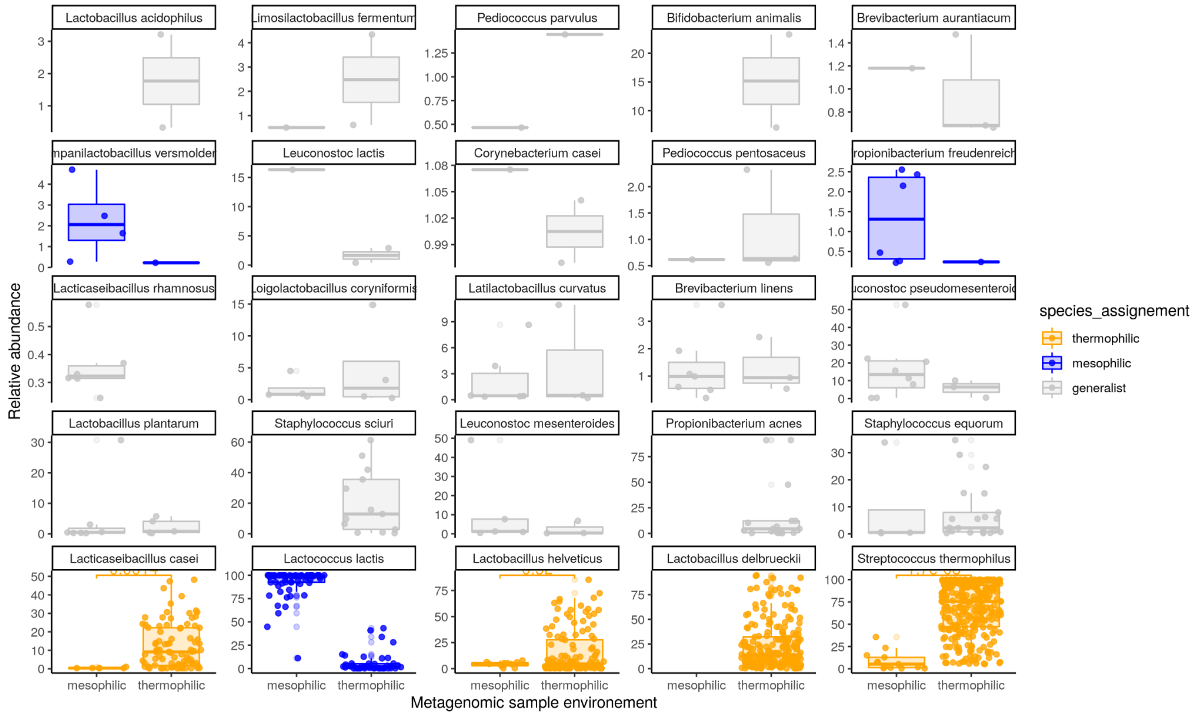


Supplementary Figure 2. Species environment assignment. The relative abundance of the dominant species in the mesophilic and thermophilic metagenomic samples (as indicated in Fig. 1A). Also the colors indicate the species assignment to either thermophilic, mesophilic or generalist species.

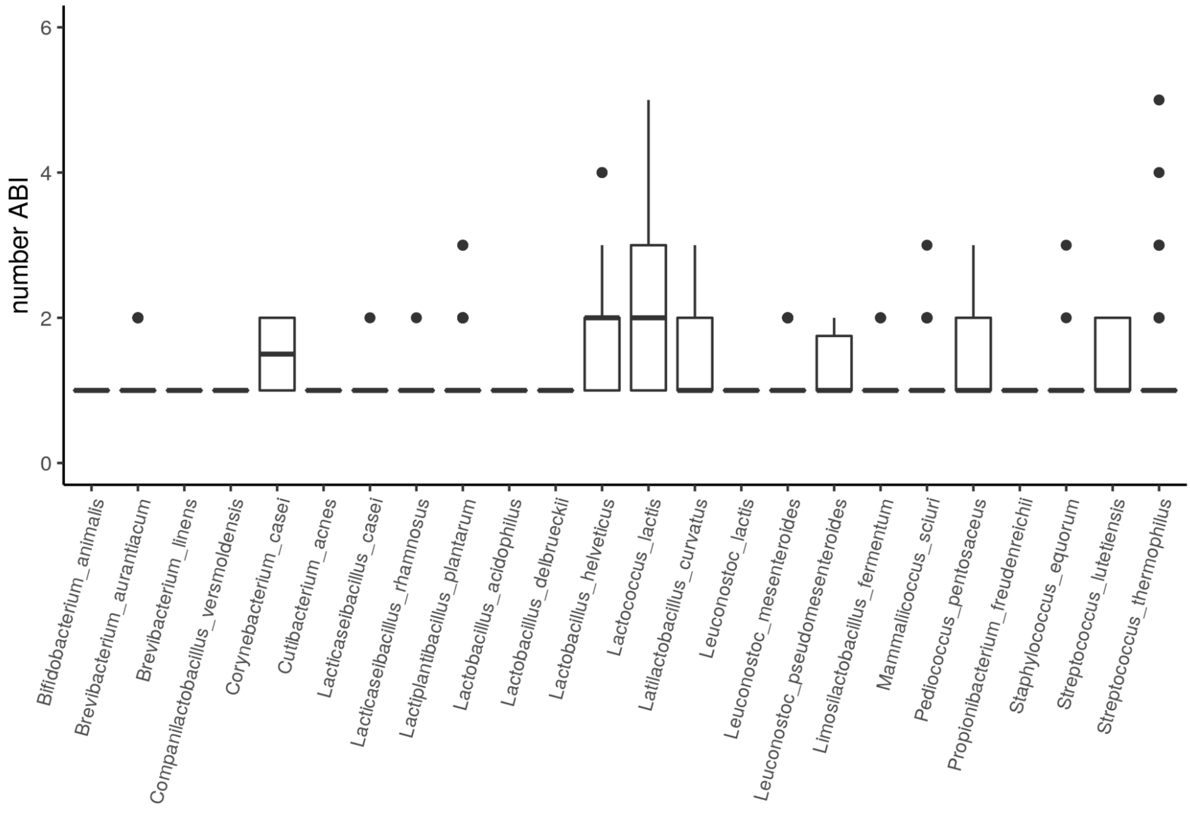


Supplementary Figure 3. Number of ABI systems per strain across different species.

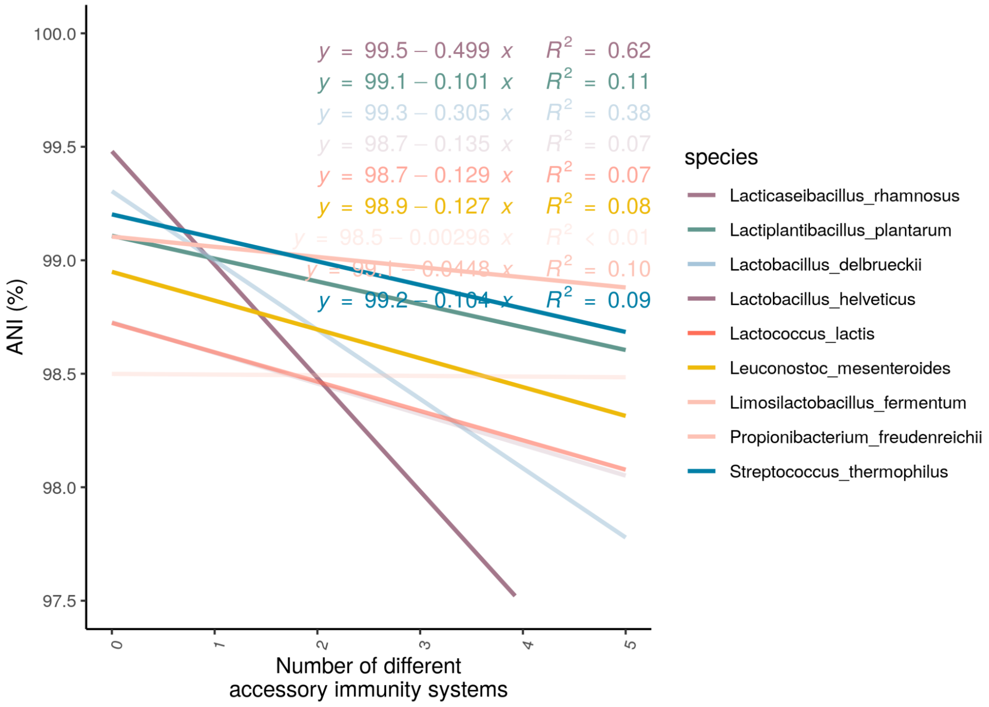


Supplementary Figure 4. The number of different defense systems vs. average nucleotides between two genomes of the same species. Including only the most dominant species comparisons. This figure corresponds to Fig. 1G in the main text but includes the regression statistics.

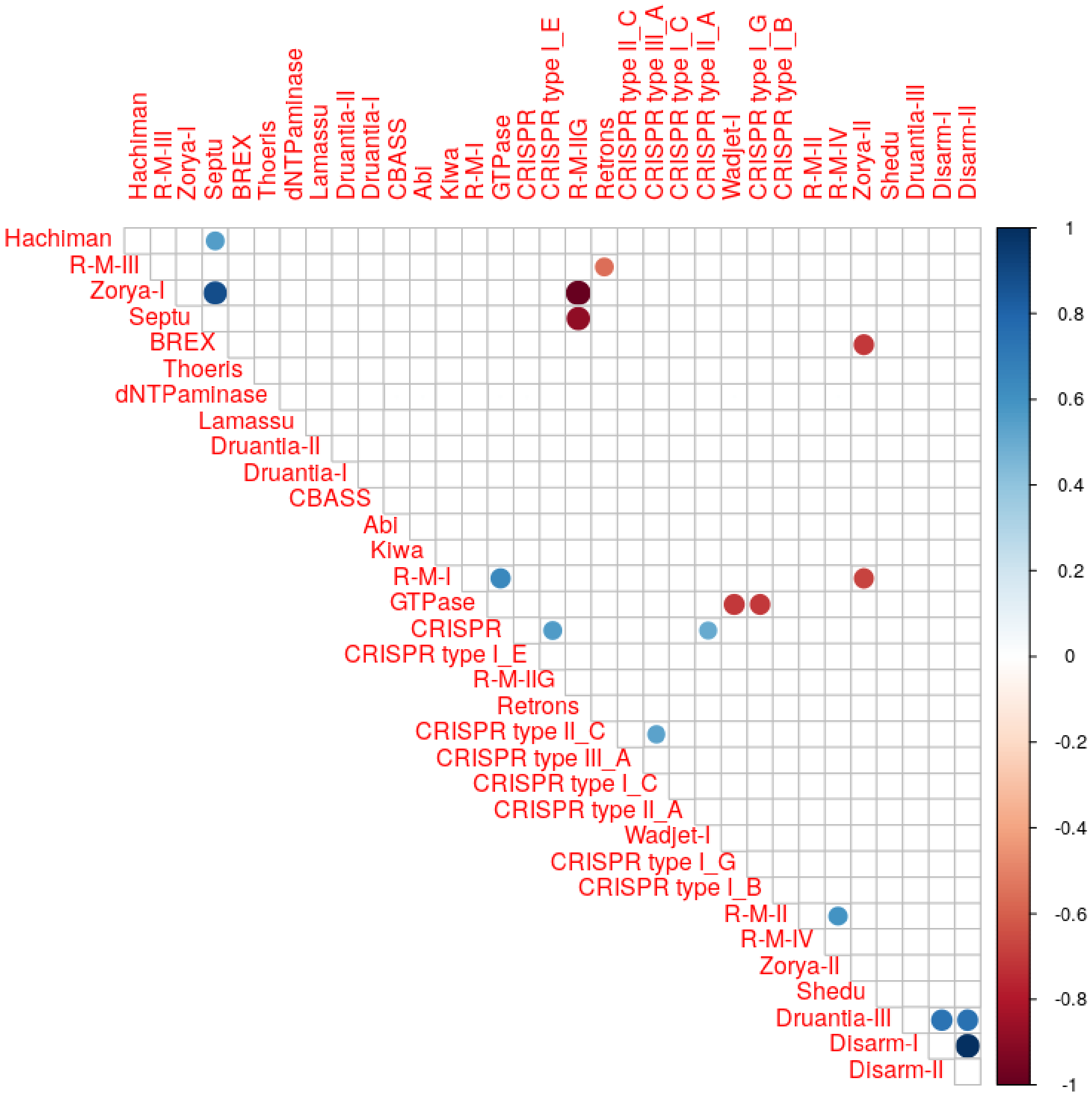


Supplementary Figure 5. The correlations between different phage defense systems. The heatmap illustrates the correlation coefficient (see legend to the right).

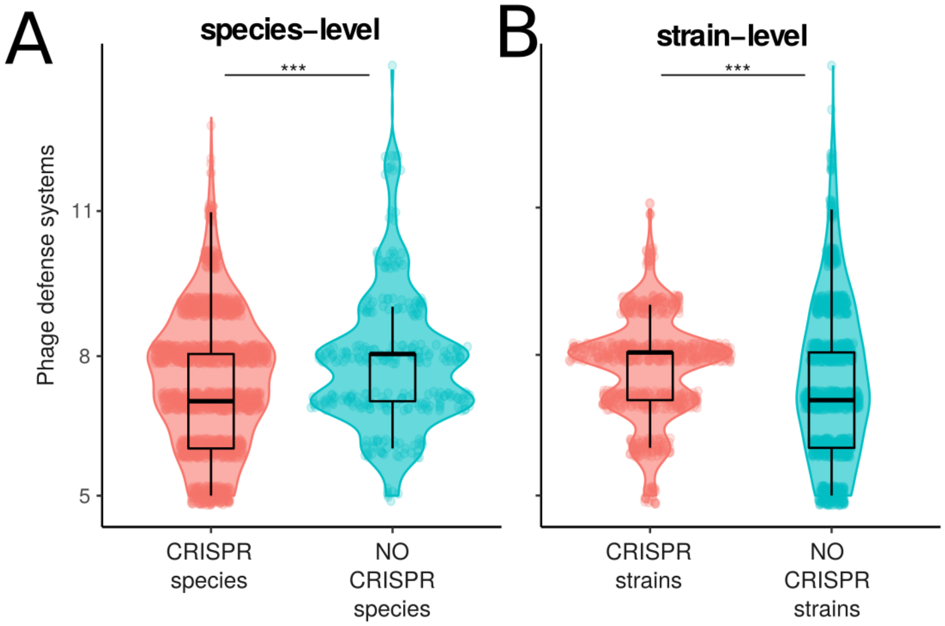


Supplementary Figure 6. Number and variation of phage defense systems between A) CRISPR containing and no CRISPR containing species and B) within the CRISPR-containing species between CRISPR containing and no CRISPR containing strains.

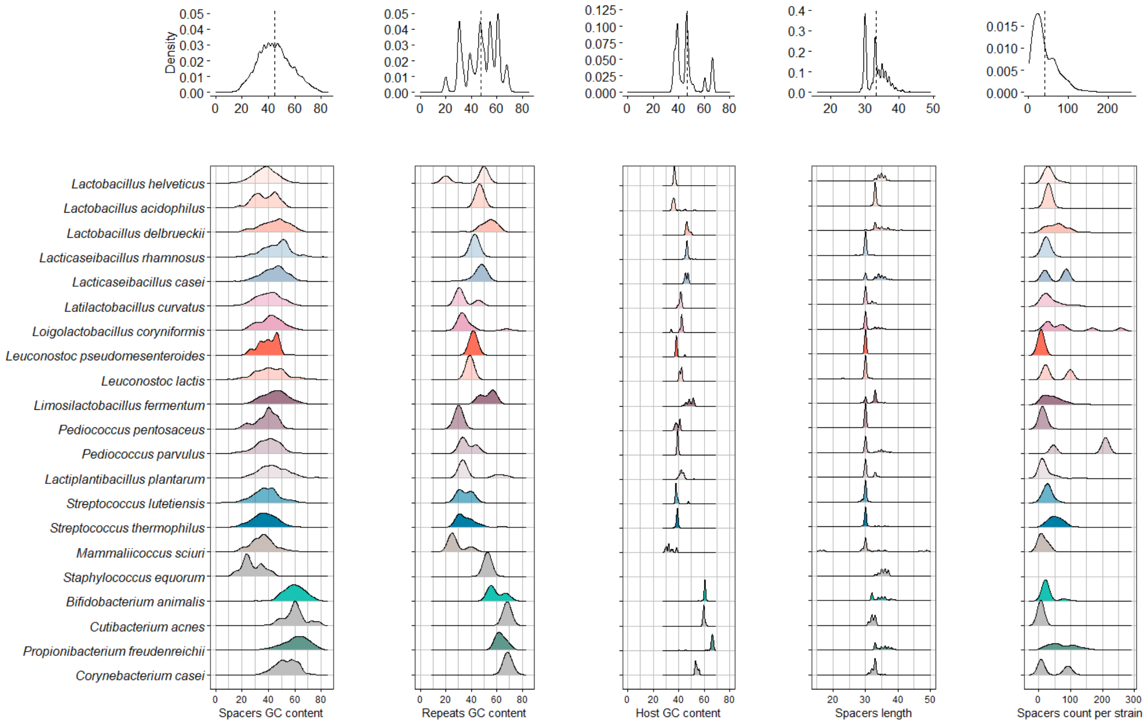


Supplementary Figure 7. Characteristics of CRISPR spacer and CRISPR repeats in the isolates of the different cheese-associated bacterial species illustrated by the following panels i) spacer GC content, ii) repeat GC content, iii) host GC content, iv) spacer length and v) number of spacers per strain.

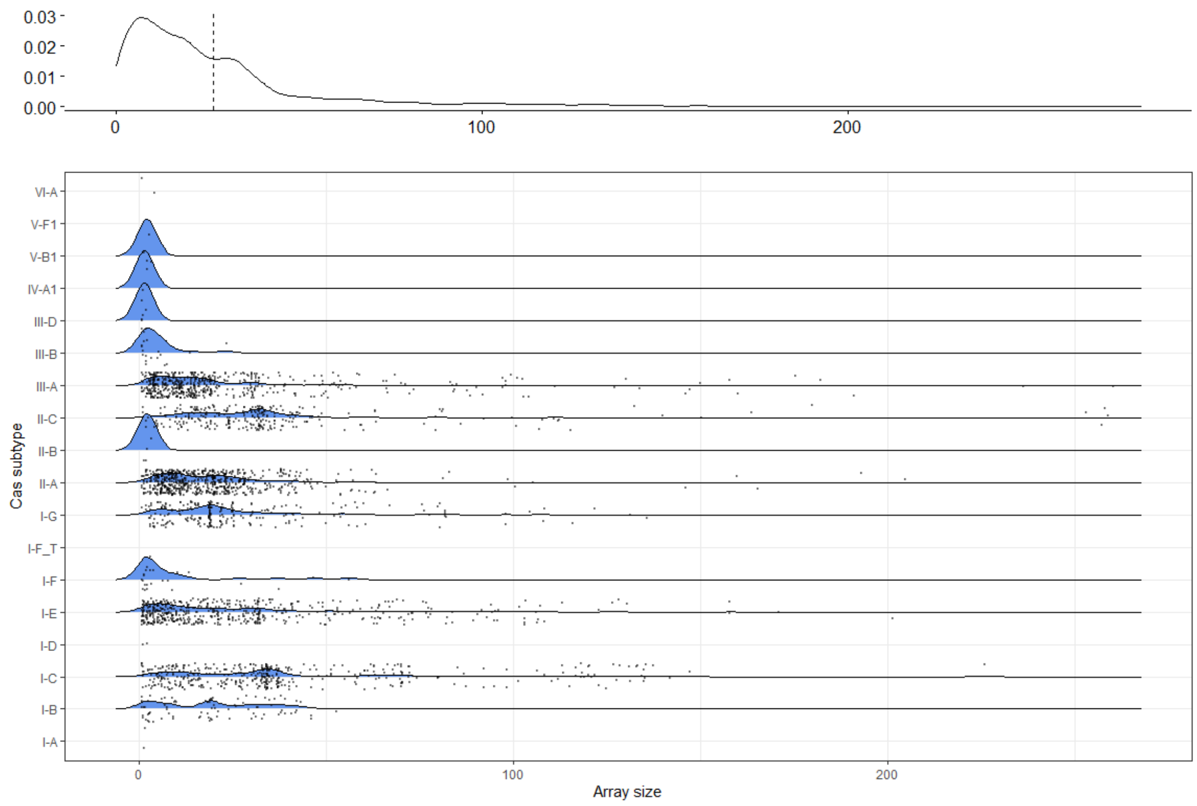


Supplementary Figure 8. The total number of CRISPR spacers per array is illustrated for Cas subtypes independent of species. Both the distribution as well as the actual number (points) are illustrated.

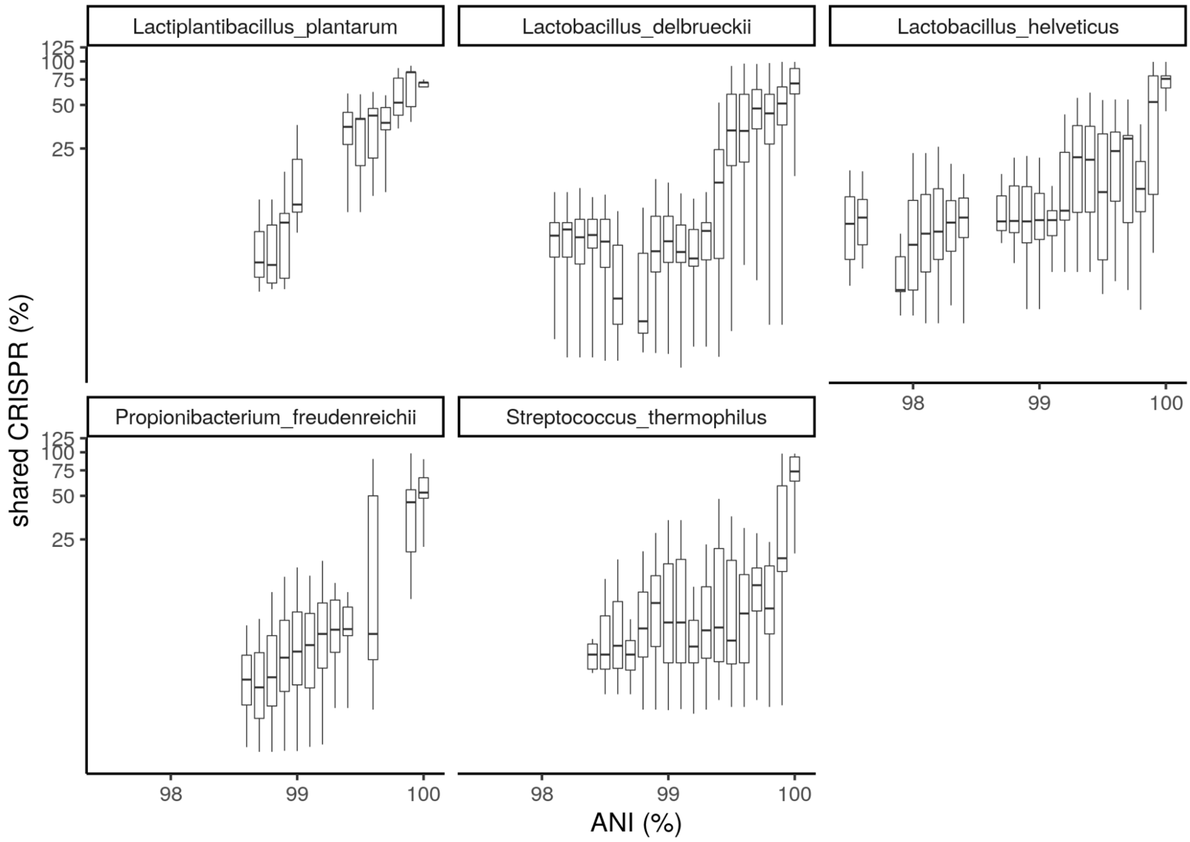


Supplementary Figure 9. The ANI vs. shared CRISPR spacer plot shown for the predominant species separately. Only boxes with more than 30 comparisons are shown.

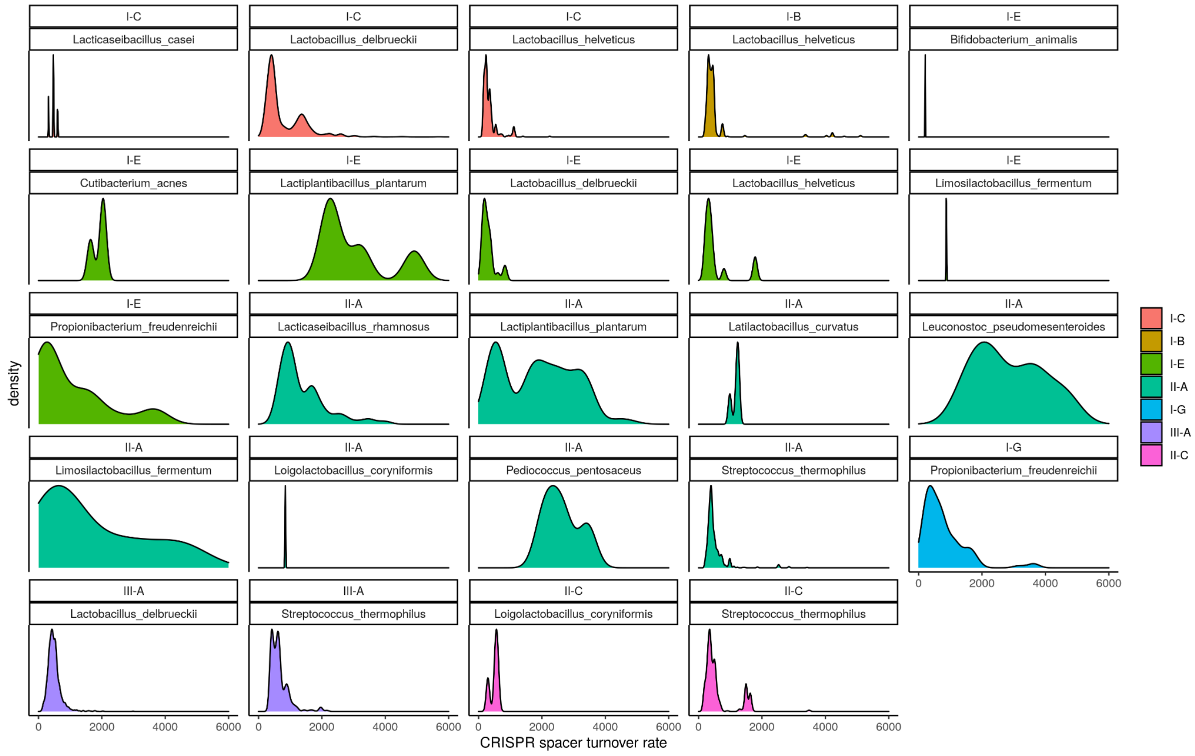

Supplementary Figure 10. CRISPR spacer turnover rates divided by species and CRISPR-cas subtype. The colours represent the CRISPR-cas subtype. Only the most dominant species are illustrated.

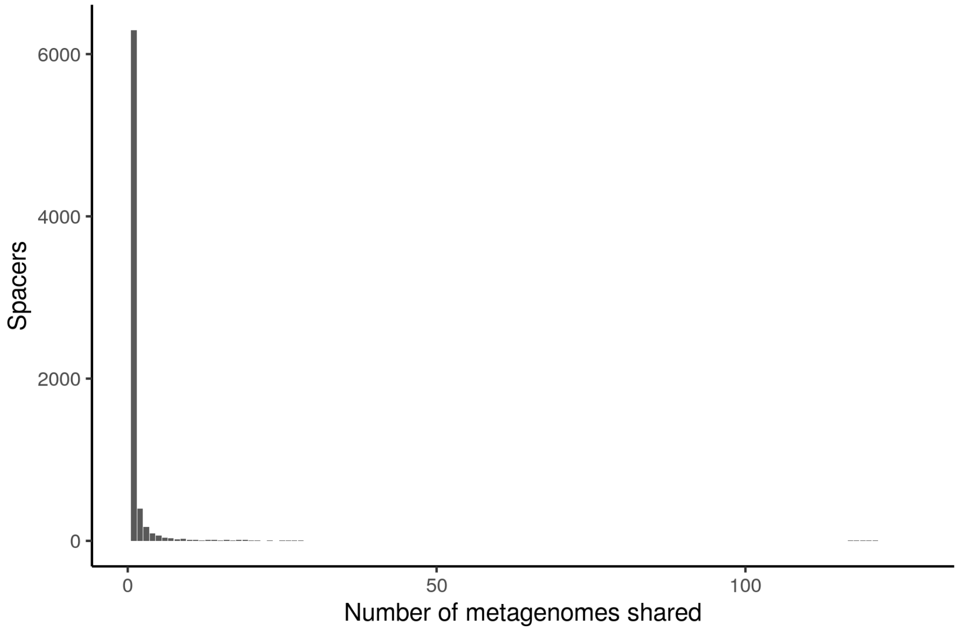


Supplementary Figure 11. The number of metagenomes that a spacer occurred in. If a spacer is unique for a single metagenome it is represented in the first column. Thereafter the columns illustrate the number of metagenomes a spacer is shared in. The large majority of spacers occur only in one or a few metagenomic samples.

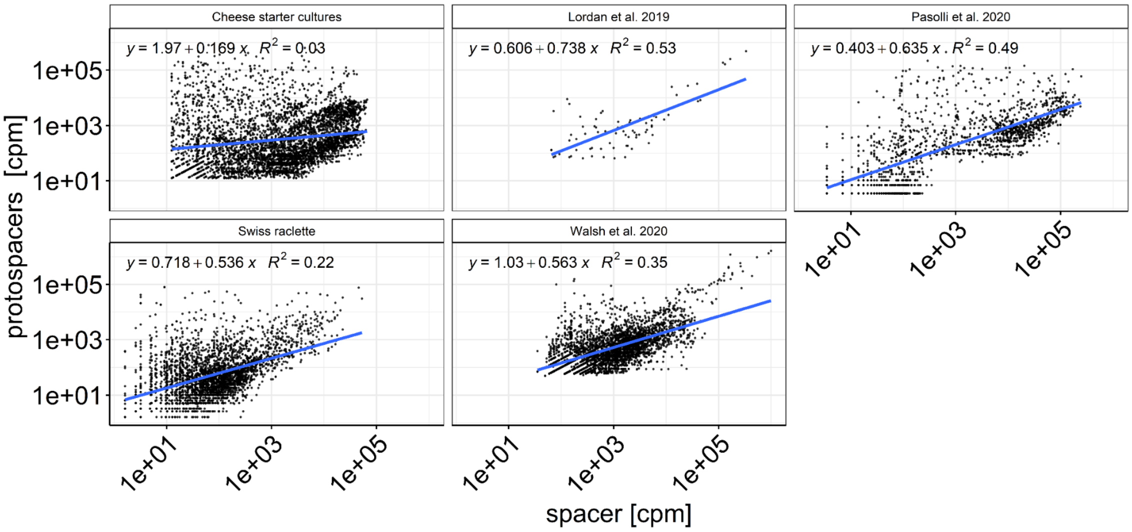


Supplementary Figure 12. Normalized spacer vs. protospacer abundances (counts per million) subdivided by the different metagenomic bioprojects. The blue line indicates the illustrated linear regression with the statistics represented at the top of the figures.

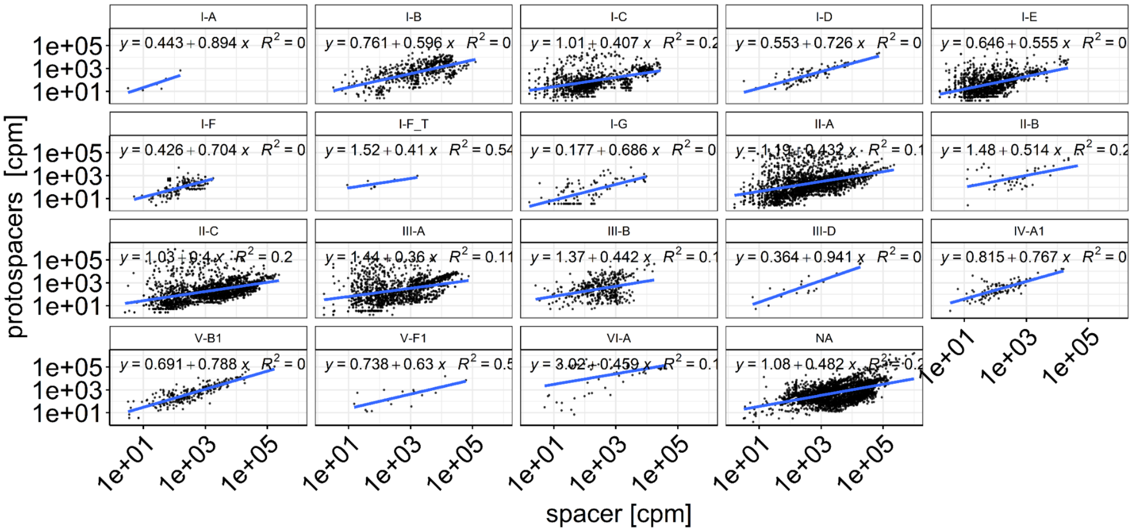


Supplementary Figure 13. Normalized spacer vs. protospacer abundances (counts per million) subdivided by CRISPR-cas subtypes. The blue line indicates the illustrated linear regression with the statistics represented at the top of the figures.

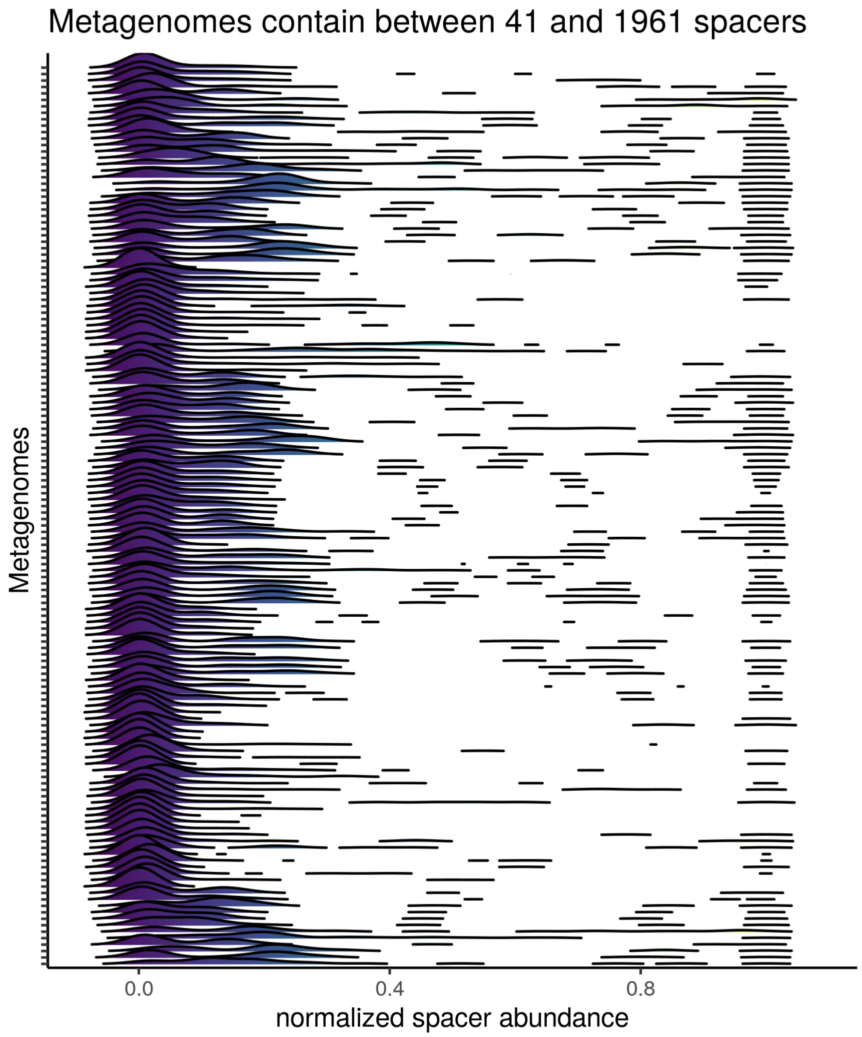


Supplementary Figure 14. The normalized spacer abundance (normalized by the most abundant spacer in the metagenome) of all metagenomic samples. Every individual metagenome contained between 41 and 1961 spacers.The large majority of spacers are of very low abundance. Apart from the accumulation of spacers at the low abundance spectrum there does not seem to be a large accumulation of spacers throughout the figure. This would have illustrated a dominant strain.

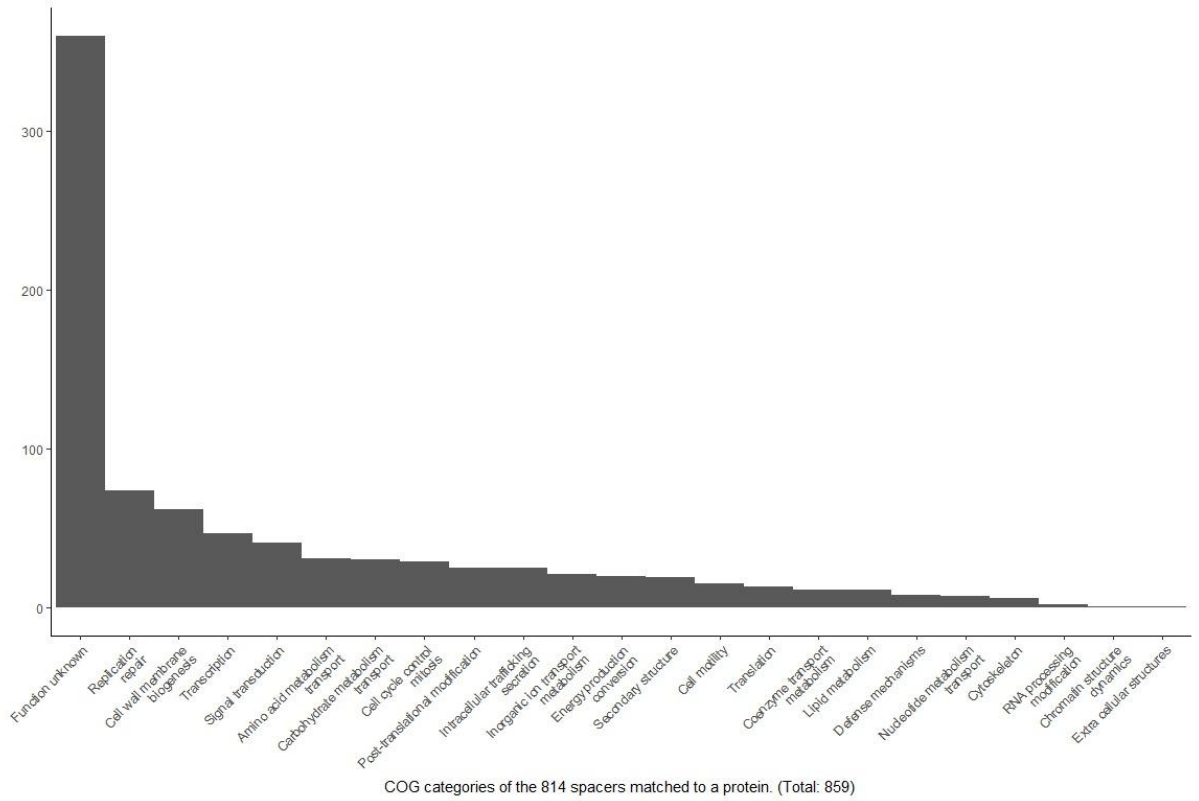


Supplementary Figure 15. COG categories of the 859 proteins that were targeted by spacers (i.e. protospacers).

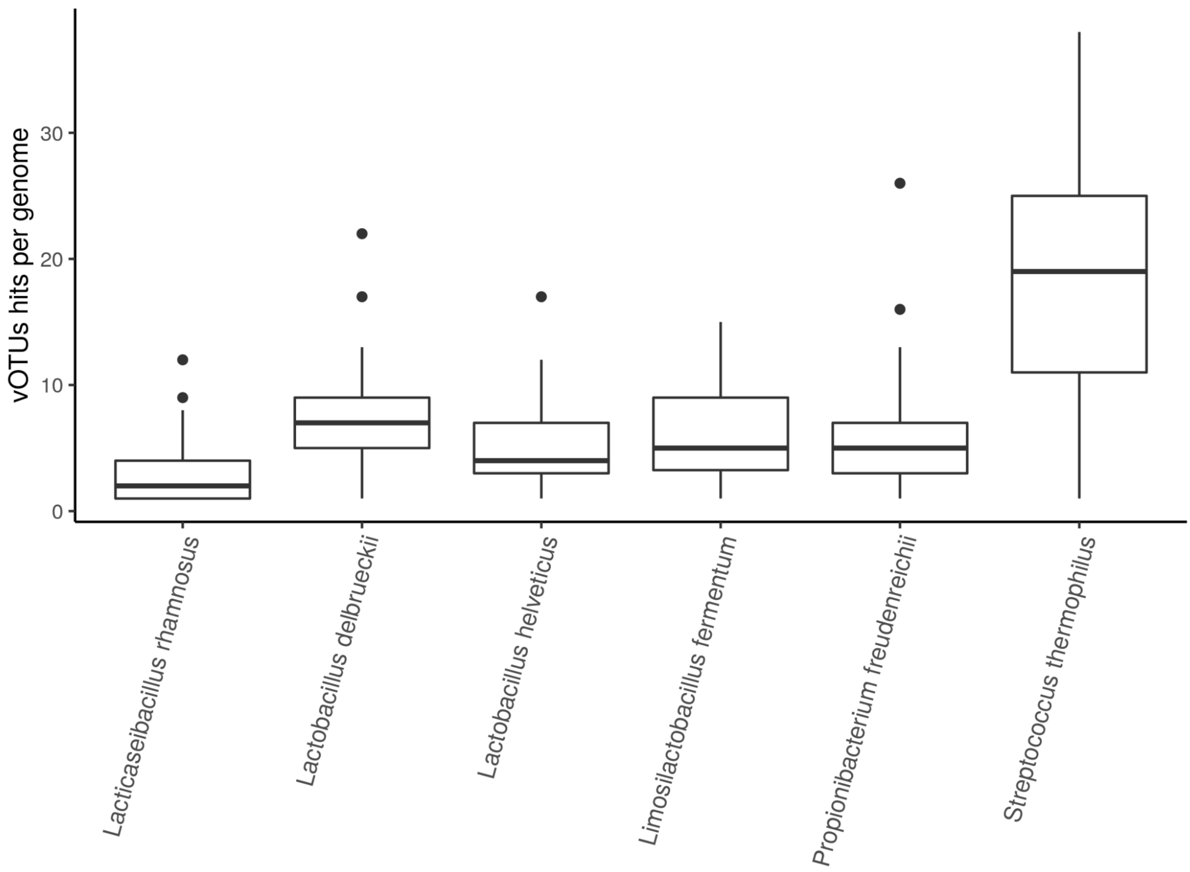


Supplementary Figure 16. The number of vOTUs targeted by a given genome of the different species. Only depicted for the most abundant species and vOTUs.

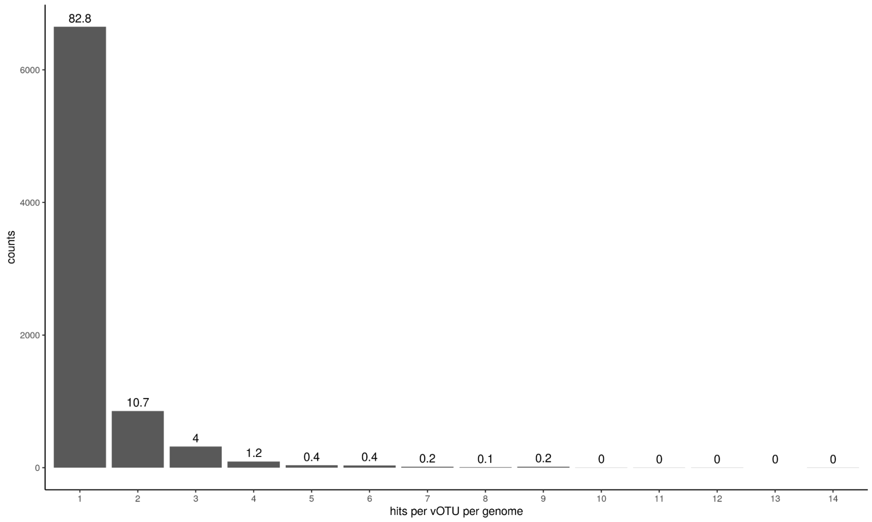


Supplementary Figure 17. The number of CRISPR spacers targeting the same vOTU in one genome. The percent is indicated above the bar. The large majority of spacers within a CRISPR array (~83%) target only a single vOTU.

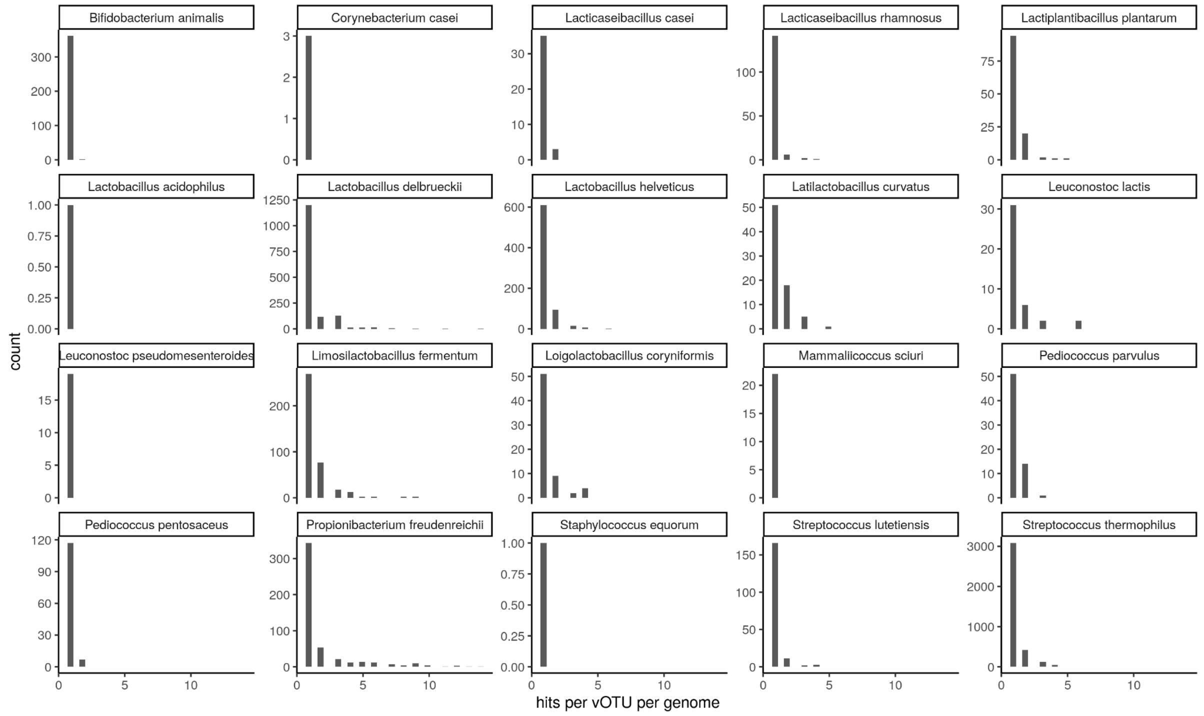


Supplementary Figure 18. Similar as the previous supplement figure, the number of CRISPR spacers targeting the same vOTU in one genome subdivided by the different species. The percent is indicated above the bar. Also here, the large majority of spacers within a CRISPR array target only a single vOTU.

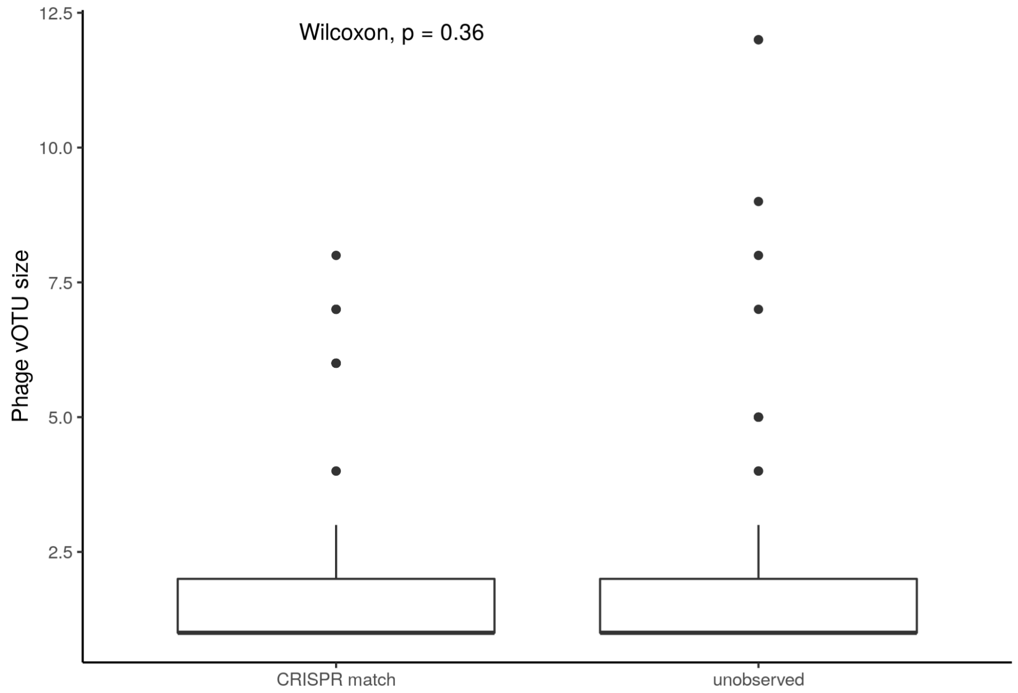

Supplementary Figure 19. The cluster size of the vOTUS (indicating the number of known/sequence phages for this vOTU) for the vOTUs which have a CRISPR match in comparison to the vOTUS that remained unobserved (no target identified in the genomes). The Wilcoxen p-value is non-significant and indicated in the figure.

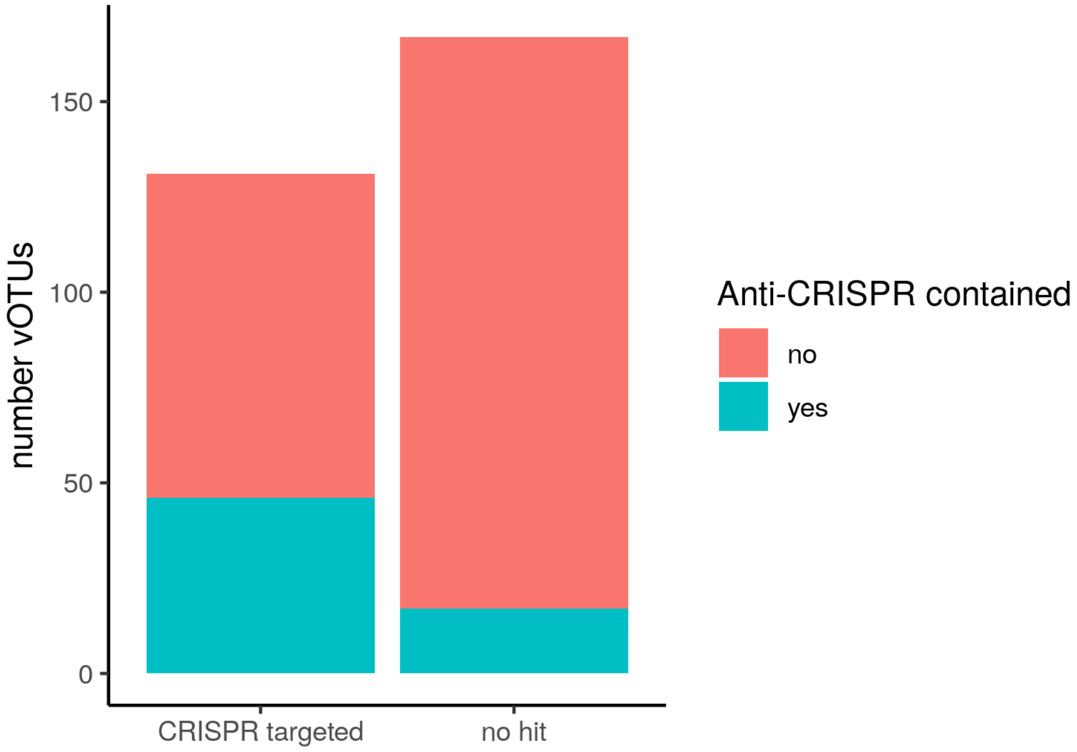


Supplementary Figure 20. The number and fraction of vOTUs containing an anti-CRISPR sequence divided by vOTUs targeted by CRISPR or not targeted (no hit) that contain anti-CRISPR genes.
